# Cell specification and functional interactions in the pig blastocyst inferred from single cell transcriptomics

**DOI:** 10.1101/2023.05.30.542847

**Authors:** Adrien Dufour, Cyril Kurilo, Jan B. Stöckl, Denis Laloë, Yoann Bailly, Patrick Manceau, Frédéric Martins, Ali G. Turhan, Stéphane Ferchaud, Bertrand Pain, Thomas Fröhlich, Sylvain Foissac, Jérôme Artus, Hervé Acloque

## Abstract

The early embryonic development of the pig comprises a long *in utero* pre- and peri-implantation development, which dramatically differs from mouse and human. During this protracted peri-implantation period, an intimate dialogue between the embryo and the uterus is established through a complex series of paracrine signals. It leads to concomitant drastic changes in the embryonic morphology and uterine receptivity to implantation. From day 7 after fertilization, the spherical blastocyst elongates as ovoid, tubular and filamentous wich transforms from a 0.5-1 mm diameter sphere to a 1000 mm long filamentous blastocyst at day 16. At the same time, the inner cell mass moves up to the trophoblast as the Rauber’s layer disappears and evolves as an embryonic disc that is directly exposed to molecules present in the uterine fluids. These drastic changes occurring before implantation are driven and coordinated by interactions between embryonic cells and maternal tissues.

To better understand the biology of pig embryos before implantation, we generated a large dataset of single-cell RNAseq at different embryonic stages (early and hatched blastocyst, spheroid and ovoid conceptus) and proteomic datasets from corresponding uterine fluids. These data were cleaned, filtered and represent a total of 34,888 cells. We first characterised the embryonic and extra-embryonic cell populations and their evolution, and identified population-specific markers of the three main populations (epiblast, trophectoderm and hypoblast). Our analysis also confirmed known functions and predicted new biological functions associated with these cell populations.We then inferred gene regulatory networks working on modules of gene regulation (regulon) and selected those that were specifically active in each embryonic population. We confirmed the relevance of the identified regulons by a meta-analysis of two other scRNAseq datasets (porcine and human preimplantation embryos). We then linked these regulons to signalling pathways and biological processes. Our results confirm the molecular specificity and functionality of the three main cell populations and identify novel stage-specific subpopulations. In particular, we discovered two previously unknown subpopulations of the trophectoderm, one characterised by the expression of LRP2, which could represent a subpopulation of progenitor cells, and the other, expressing many pro-apoptotic markers, which could correspond to the cells of the Rauber’s layer. We also provide new insights into the biology of these populations, their reciprocal functional interactions and the molecular dialogue established with the maternal uterine environment.

## Introduction

The pig is a species of increasing interest both as a biomedical model for human pathologies [1], as one of the main source of animal protein for human nutrition and as an alternative to rodent animal models for the study of early mammalian development [2]. The early embryonic development of the pig comprises a long *in utero* pre- and peri-implantation process, which dramatically differs from mouse and human. During this protracted peri-implantation period, an intimate dialogue between the embryo and the uterus is established through complex series of paracrine and exocrine signals [2]. This leads to concomitant changes in the uterine receptivity to implantation and the embryonic morphology. After fertilisation, the porcine embryo undergoes a series of cleavages. At 4 days post-fertilisation (dpf), the embryo undergoes the process of compaction, which is associated with an increase in intercellular adhesion and the acquisition of cell polarity, giving it the appearance of a mulberry, named morula. At 5 dpf, it undergoes the second major morphogenetic event which is the formation of the blastocyst, characterised by a fluid-filled cavity known as the blastocoel. This process is closely linked to the specification of the inner cell mass (ICM) and the trophectoderm (TE). At 6 dpf, the hypoblast (HYPO or primitive endoderm) and the epiblast (EPI) are specified from the ICM. From 7 dpf, the spherical blastocyst elongates as ovoid, tubular and filamentous blastocyst, transforming from a 0.5-1 mm diameter sphere to a 1000 mm long filamentous blastocyst at 16 dpf. At the same time, the inner cell mass moves up to the trophectoderm (TE) as the polar TE (Rauber’s layer) disappears and forms the epiblast (EPI). It then develops as an embryonic disc directly exposed to molecules present in the uterine fluids. These drastic changes, which occur before implantation, are likely controlled and coordinated by key functional interactions between cells and tissues.

Until recently, the study and interpretation of these interactions has been hampered by the lack of molecular tools and datasets that allow the implementation of systems biology approaches. Most of the knowledge about these interactions was inferred from observations made in other mammalian species, in particular mice, humans or cattle [3]–[5]. Classical genes and pathways known to control early embryonic and extra-embryonic cell specification have been tested in cultured pig embryos or explants [6]–[9], but it has been difficult to gain a deeper understanding of such mechanisms due to the difficulty of accessing pig embryos at late stages of pre-implantation development, to the lack of *bona fide* embryonic and extra-embryonic stem cells and to the difficulty to perform functional genomics.

The recent development of single-cell RNA sequencing (scRNAseq) has enabled remarkable advances in the understanding of the first steps of mammalian embryogenesis, from the definition of embryonic and extra-embryonic populations to the gene regulatory networks controlling cell fates. Indeed, scRNAseq studies of mouse pre-implantation embryos highlight the switch from the naive pluripotent state observed in the ICM of the blastocyst to the formative and primed pluripotent states observed in the epiblast, coinciding with the sequential specification of the trophectoderm and the hypoblast cell populations [3]. Recently, scRNAseq studies of porcine embryos have also been published, describing the switch from a naive epiblast (from 4dpf to 6dpf) to a primed epiblast (from 7dpf) [10], [11], which is associated with a switch in signalling pathways. In the ICM, the IL6-STAT3 and PI3K-AKT pathways are mainly activated and regulate the expression of markers of naive embryonic pluripotency (KLF4, ESRRB, STAT3). When the EPI forms, these two signalling pathways decrease in favour of the TGFβ-SMAD2/3 pathway, which in turn regulates the expression of markers of primed pluripotency (NANOG, DNMT3B, OTX2) and is associated with a metabolic switch between OXPHOS and glycolysis. From 10dpf, these studies described a primed state of pluripotency in the epiblast where canonical Wnt signalling activity increases and primes the pluripotent epiblast for gastrulation and mesendoderm formation [10], [11]. These studies also suggest a potential conserved role of IL1B genes between pig and human/monkey for implantation and rapid trophectoderm expansion in pig. However, as these studies have mainly focused on EPI, information on other extra-embryonic cell populations is still lacking. This is mainly due to the difficulty to obtain a large number of cells representative of all embryonic and extra-embryonic subpopulations using the smart-seq2 approach [12]. Here we used the Chromium 10x Genomics technology to provide a scRNAseq analysis of pig embryos at four different embryonic stages: (1) early blastocyst (E5), (2) hatched blastocyst (E7), (3) spheroid/early ovoid blastocyst (E9) and (4) late ovoid blastocyst (E11). Proteomic datasets were also generated from the uterine fluids of the sows used for embryo production. We characterised a panel of 34,888 cells, from which we first characterised embryonic and extra-embryonic cell populations and their evolution, and identified population-specific markers of the three main populations (epiblast, trophectoderm and hypoblast). We also identified known and novel specific functions associated with the biology of these subpopulations. We then inferred gene regulatory networks working on modules of gene regulation (regulon) and selected those that were specifically active in each embryonic population. We then linked these regulons to signalling pathways and biological processes. To do this, we constructed signalling networks from ligands (expressed by cells or present in the uterine fluids), receptors, intermediaries and transcription factors. Our results confirm the molecular specificity and functionality of the three main cell populations and identify novel stage-specific subpopulations. We also provide new insights into the biology of these populations, into their reciprocal functional interactions and the molecular dialogue established with the maternal organism through the uterine fluids.

## Results and discussion

### 1. Identification of embryonic and extra-embryonic cell populations and their associated biological functions

Using the Chromium 10x Genomics technology, we generated a large dataset of scRNAseq at four different embryonic stages corresponding to: (1) early blastocyst (E5), (2) hatched blastocyst (E7), (3) spheroid/early ovoid blastocyst (E9) and (4) late ovoid blastocyst (E11) (Figure 1A and supplementary Table 1). Raw reads were mapped, re-attributed to each cell and counted. A minimum number of UMIs per cell, a maximum percentage of mitochondrial transcripts per cell and a maximum number of features per cell were used to exclude the cells that did not reach a sufficient quality level (supplementary Table 2). We validated 34,888 cells, distributed as follows: 1,226 cells at E5, 4,228 cells at E7, 12,727 cells at E9 and 16,707 cells at E11 (Figure 1B and supplementary Table 2).

**Figure 1.**
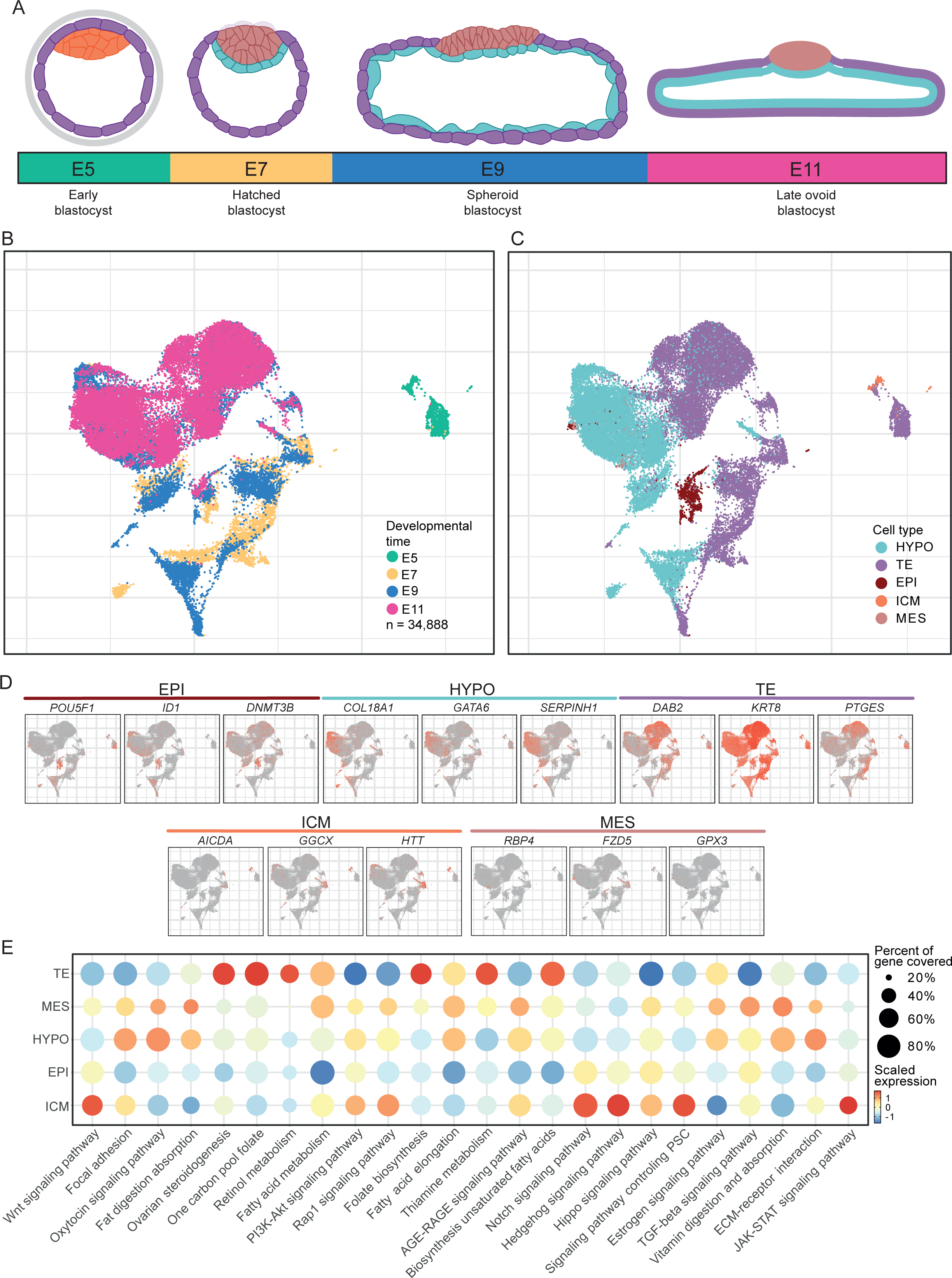
Dynamic evolution of cell lineages from the early to the ovoid blastocyst stages (E5 to E11) (A) Schematic view of pig embryo morphology from embryonic day (E) 5 to E11. Cells from the inner cell mass and the epiblast are represented in orange/red, the trophectoderm in purple and the hypoblast in turquoise. (B) Visualization of cells coloured by developmental time via UMAP: E5 (green), E7(yellow), E9 (blue) and E11 (pink). (C) Identification of five major clusters coloured by population via UMAP: inner cell mass (ICM, orange), epiblast (EPI, red), hypoblast (HYPO, turquoise), trophectoderm (TE, purple) and mesendoderm (MES, light red). (D) UMAP plot of gene markers for each population. (E) Dot plot visualization of selected KEGG signalling pathways. The circle size represents the percentage of genes out of all the genes in the pathways that are expressed by the cell populations. The red gradient represents the mean scaled expression of the pathways within the cell populations. AGE-RAGE signalling pathway in diabetic complications has been abbreviated to AGE-RAGE signalling pathway.

From this dataset, to identify transcriptionally distinct cell populations, we performed a dimensionality reduction and clustering approach using Principal Component Analysis (PCA) and Uniform Manifold Approximation and Projection (UMAP) following the Harmony and Seurat workflows. To identify the distinct cell populations, we then visualised known population marker genes from the literature for ICM (*AICDA*, *GGCX, HTT*), EPI (*POU5F1*, *ID1*, *DNMT3B*), TE (*DAB2, KRT8, PTGES*), HYPO (*COL18A1*, *GATA6*, *SERPINH1*) and putative early embryonic mesendoderm (*RBP4*, *FZD5, GPX3*) (Figure 1C and 1D).

We then searched for enriched functions in these five cell populations by performing a gene set variation analysis (GSVA) (Supplementary Table 3c) using the expressed genes within each lineage. The most significantly enriched pathways and those selected from the literature are shown in Figure 1E. We found that the pool of genes corresponding to Notch, JAK-STAT, Hippo, Hedgehog and Wnt signalling pathways was enriched in ICM cells compared to other cell populations but similar pathways (except JAK-STAT) were also enriched in EPI. In TE, we found an enrichment of genes associated with ovarian steroidogenesis and estrogen signalling pathway, fatty acid elongation and unsaturated fatty acids and folate biosynthesis, one-carbon, retinol and thiamine metabolism, reflecting known TE biological functions. In the HYPO, we observed an enrichment of genes associated with the oxytocin signalling pathway, fat digestion and absorption, fatty acid elongation, vitamin digestion and absorption, focal adhesion and ECM-receptor interaction.

We next used SCENIC to identify regulons, which are defined as functional modules of gene regulation. Each regulon associates a transcription factor (TF) and its direct target genes, which are defined by their co-expression with the TF and by sharing a common binding motif for this TF in their promoters [13]. For each regulon, activity scores were calculated for each cell using the relative expression of the gene that makes up the regulon. 297 regulons were identified and their activity score was summarised by their mean expression in each cluster at each state (Supplementary Table 4a). In parallel, we performed the same analysis on two publicly available scRNAseq datasets from pig and human pre-implantation embryos [3], [11] and we looked for common regulons across these scRNAseq analyses (Supplementary Figure 1). We then generated a selection of the most specific regulons for each cell population based on the Regulon Specificity Score (RSS) (Supplementary Table 4b).

### 2. Timely diversification of TE cell population with distinct molecular functions

To further characterise the TE cell population, we selected 18,239 cells from our dataset corresponding to the TE population and performed dimensionality reduction followed by a new clustering (Figure 2A). This led to the identification of eight TE subpopulations (Figure 2B). All of these subpopulations show a clear TE signature by expressing *GATA2, GATA3, DAB2* and *PTGES* (Figure 2C), but each has specific characteristics.

**Figure 2.**
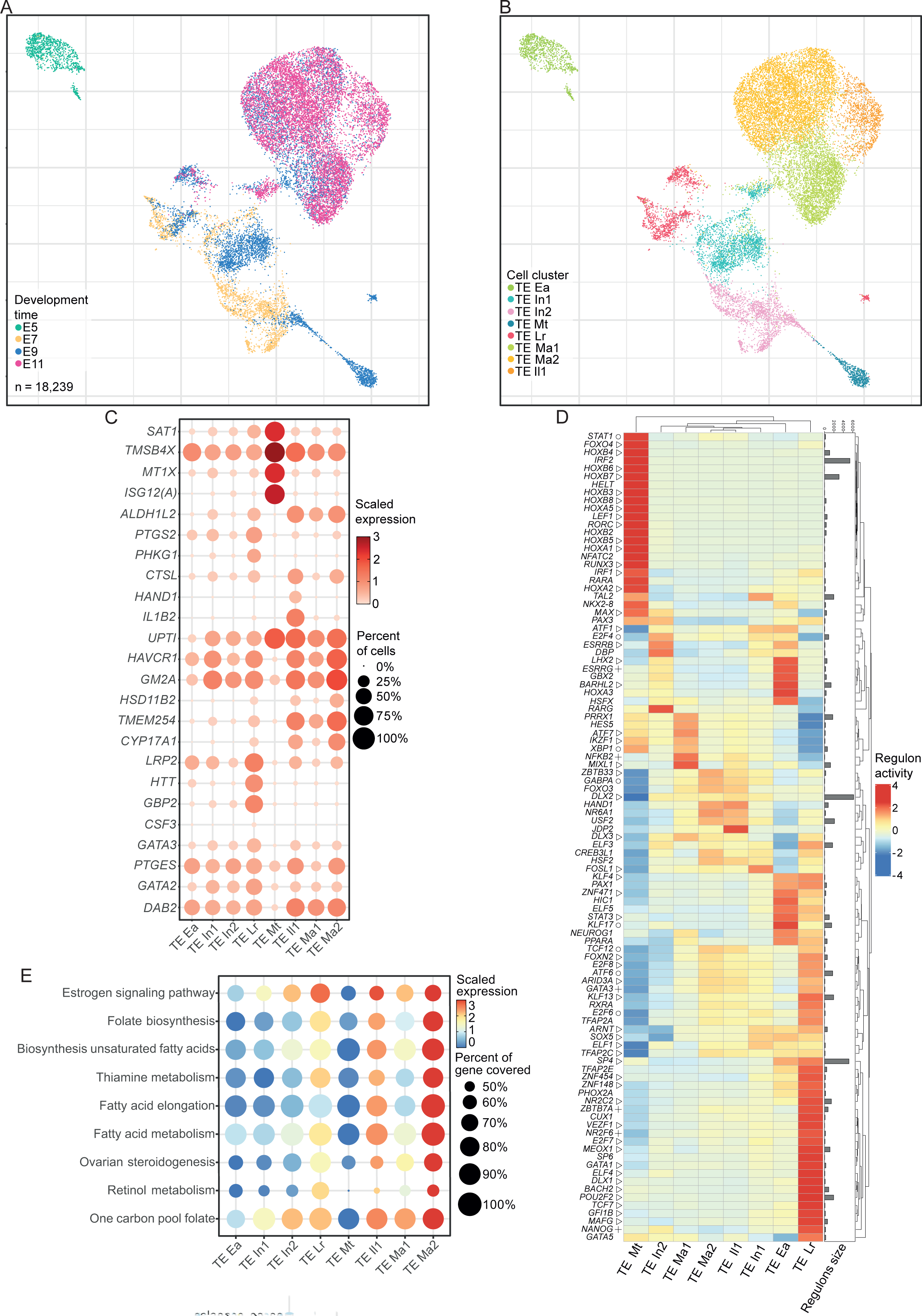
Identification and characterisation of different populations in the trophectoderm. (A) Visualisation of selected TE cells coloured by developmental day via UMAP: E5 (green), E7 (yellow), E9 (blue), E11 (pink). (B) Visualisation of TE populations coloured by cluster via UMAP: TE Ea (green), TE In1 (light blue), TE In2 (pink), TE Lr (red), TE Mt (dark blue), TE Ma1 (green), TE Ma2 (yellow), TE Il1 (orange). (C) Dot plot visualization of selected DEG genes, the circle size represents the percentage of cells within the cluster that express the gene. The red gradient represents the mean scaled expression of the genes within the cluster. (D) Heatmap showing scaled values of Regulon Activity Score for the 20 most specific regulons for each cluster, identified by Regulon Specificity Score (RSS). (△): common regulons with another pig study (+): common regulons with another human study; (Օ): common regulons within the three studies. Right rows (heatmap): histograms distribution of regulon size (number of genes regulated by the TF in the regulons. (E) Dot plot visualization of selected KEGG signalling pathways, the circle size represents the percentage of genes within all the genes in the pathways that express by the clusters. The red gradient represents the mean scaled expression of the pathways within the cluster.

At the early blastocyst stage (E5), the first cell lineage decision leads to the formation of the early trophectoderm (TE Ea, light green dots in Figure 2B). This subpopulation is characterised by the expression of early TE marker genes such as *DAB2*, *GATA2* and *PTGES* [11], [14], [15] (Figure 2A and Supplementary Table 3a). Early TE appeared to be quite distinct from other TE subpopulations in terms of gene expression (Figure 2B, Supplementary Table 3a), with functional enrichment of genes related to aerobic respiration and cell metabolism (Supplementary Table 3d). We also highlighted specifically active regulons at this stage: *LHX2* and *BARHL2* (identified in our study and from data in [11]), *ESRRG* (in our study and from data in [3]), and *GBX2* and *HSFX* (our study) (Figure 2D). We also observed active *TFAP2C* and *TFAP2E* regulons, which TFs are known to be involved in ICM/TE segregation in mice [16].

Then, at the subsequent E7 hatched blastocyst stage, E9 early ovoid and E11 late ovoid stages, we identified two subpopulations that are observed at these three stages: intermediate trophectoderm 1 (TE In1, turquoise blue dots in Figure 2B) a subpopulation that expresses classical TE markers (e.g., *DAB2*, *GATA2*, *PTGES* and *GATA3*) (Figure 2A and Supplementary Table 3a) [11], [17] and another small TE subpopulation that emerges at E7 and expands in subsequent stages, which we named LRP2 TE (TE Lr, red dots in Figure 2B) due to its high expression of *LRP2, HTT* and *GBP2* (Figure 2C, Supplementary Figure 4). At E7 and E9, we also identified another subpopulation, which we termed intermediate trophectoderm 2 (TE In2). Gene expression profiles were highly similar between TE In1 and TE In2, with notable differences in mitochondrial and ribosomal genes (supplementary Table 3a), suggesting differences in their cellular metabolism: TE In2 relies on OXPHOS (together with TE Ea and TE Lr), whereas TE In1 relies on glycolysis (together with more mature TE subpopulations) (Supplementary Figure 2). Slight differences in regulon activity were also observed (Figure 2D): *ARNT* and *SOX5* regulons (identified in our study and from data in [11]) are less active in TE In2 compared to TE In1, whereas *ESRRB*, *RARA/G*, *E2F4* [20] and *DBP* are more active in TE In2. *ESRRB* is known to play a role in the transformation of stem cells into TE [18] and *RARA/G* to be involved in cell reprogramming [19]. Regulons specific to TE In1 include TFs described in cell survival such as *ATF1* [21], differentiation towards TE for *FOSL1* [22] and an unknown role for *TAL*2.

In contrast, TE Lr stands out from the other subpopulations and is characterised by several differentially expressed genes (DEG), including *IGFR*, which controls proliferation, differentiation, growth and cell survival [18], *CSF3*, which improves embryonic pig development [19] and *GBP2*, *HTT* and *LRP2* (Figure 2C, Figure 5B, Supplementary Figure 4, Supplementary Table 3a). Cells from TE Lr are cycling with more than 50% of the cells in S and G2/M phases (Supplementary Figure 3) and rely on OXPHOS metabolism (together with TE Ea and TE In2) (Supplementary Figure 2). It also presents a large set of specific regulons often associated with a stem cell signature, some of which were also detected in the ICM (*TCF7*) or in the EPI (*NANOG*), which could reflect some cell fate plasticity. It also includes *ARID3A* which has been described to be required for TE cell maintenance [16], and known TE regulators (i.e. *GATA* and *ELF* related factors, [20]) (Figure 2D). Taken together, our data suggest that TE Lr may be a population of TE progenitors that emerges around E7 and is maintained until at least E11.

**Figure 5.**
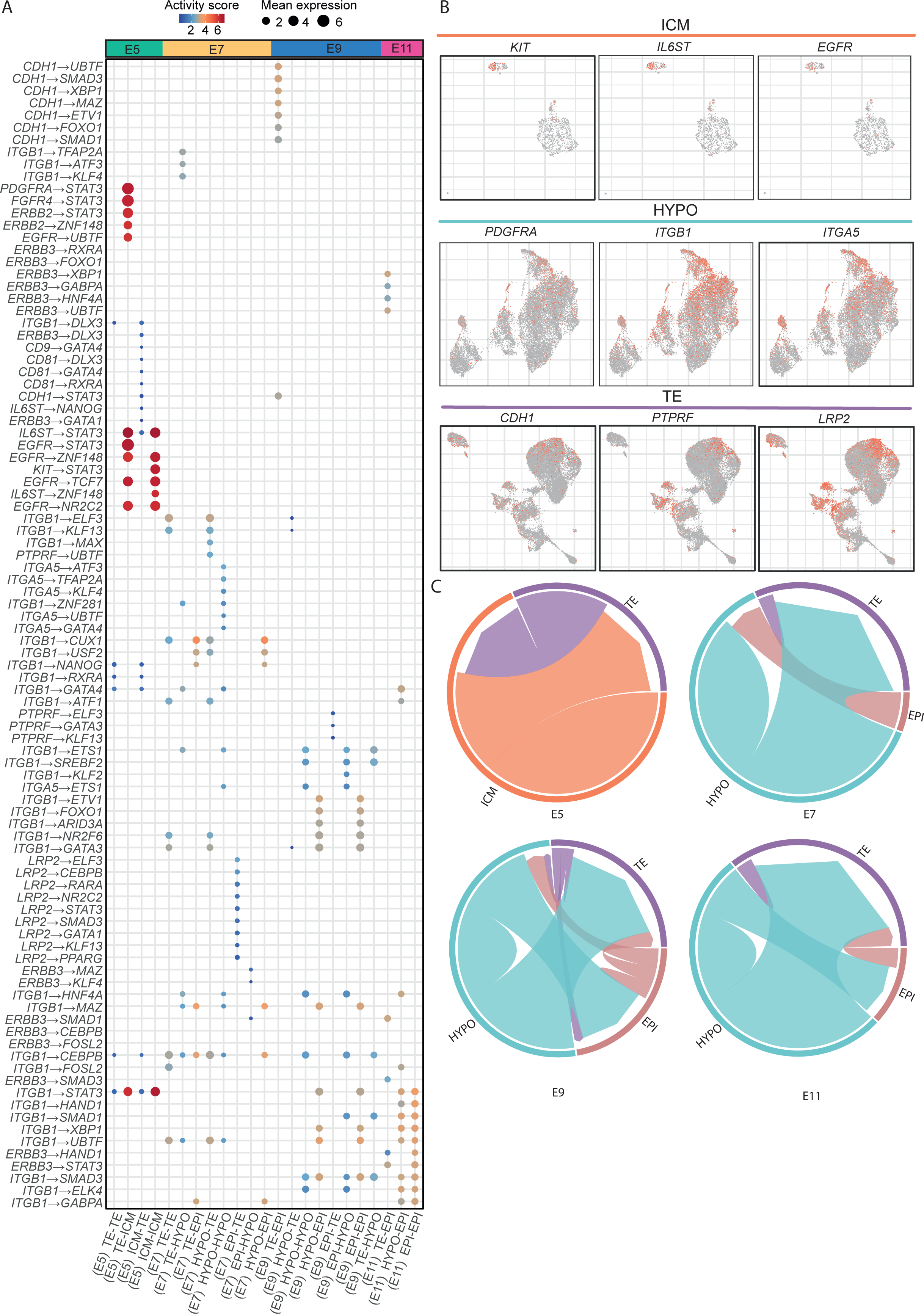
Cellular crosstalks between cell lineages from E5 to E11. (A) Dot plot displaying receptor-TF pairs identified for each cluster/cluster interaction coloured by activity score and sized by the cluster mean expression of the receptor. (B) UMAP plot of selected ligand and receptor for the ICM, HYPO and TE. (C) Chord diagram showing cellular interactions between clusters. Arrow origins represent the sum of the mean for expressed ligands and arrow ends represent the sum of the mean of expressed receptors.

Strikingly, we also observed a new TE cell population at E9, which we named Metallothionein Trophectoderm (TE Mt, blue dots in Figure 2B). This TE Mt population is only detected at E9, and its expression profile shows specific DEGs (Figure 2C and supplementary Figure 4). It is characterised by an increased expression of the metallothionein-related gene family (*MT1X*, *MT1A*), but also of *SAT1,* a gene described in human trophoblastic cell apoptosis [21], and *ISG12(A),* also known to have pro-apoptotic activity in human cells [22] (Supplementary Figure 4). The TE Mt regulons *IRF2* and *SAT1* have been described to have joint promoters with *ISG12(A)* [23], [24]. This population also shows a specific enrichment for regulons of the *HOXB* gene family, *RORC* and *HELT* (Figure 2D). We did not observe this cluster at later stages, consistent with the fact that these cells are not cycling (Supplementary Figure 3) and appear to be entering apoptosis as they strongly express pro-apoptotic genes. Taken together, this suggests that TE Mt cells may correspond to the Rauber’s layer that disappear around these stages [25].

At the ovoid stage (E9 and E11), we inferred three TE subpopulations that are specific of this stage and characterized by a more mature state of differentiation: Mature1 TE (TE Ma1, light green dots in Figure 2B), Mature2 TE (TE Ma2, yellow dots in Figure 2B) and Interleukin-1 TE (TE Il1, orange dots in Figure 2B). These three populations are quite similar regarding gene expression with shared expression of TE markers and TE differentiation and functions (*ALDH1L2, TMEM254, CYP17A1, CTSL*, Figure 2C, Supplementary Figure 4), some of these genes (e.g., *CYP17A1*, *TMEM254*, *HSD11B2*, *GM2A* and *HAVCR1*, Supplementary Figure 4) being described to play a role in elongation [26]–[28]. TE Il1 is characterised by its elevated expression level of *IL1B2* and of various RNA coding for interleukin beta-like (e.g., *ENSSSCG00000008088*, *ENSSSCG00000033667* and *ENSSSCG00000039214,* Supplementary Figure 4). These genes have been described to be necessary for the rapid elongation of the porcine conceptus [29]. Our results support that only a subset of TE cells are expressing *IL1B* at the ovoid stage and this may be driven by specific regulons. For instance, *JDP2* displays a high activity in TE Il1 cells only, and can act together with *HAND1,* which is also highly active in this population (Figure 2D) and has been described to play a role in differentiation into giant trophoblastic cells [30]. Commonly expressed genes between TE Il1 and TE Ma include *CTSL* and *PTGS2*, that have also been found into extracellular vesicles extracted from the uterine fluid of pregnant ewes [31], suggesting a TE origin of these secreted proteins in the uterine fluids. Other DEGs (e.g., *TMSB4X*, *ALDH1L2* and *UPTI, S*upplementary Table 3a, Supplementary Figure 4) have been described in TE during placentation and conceptus elongation [32]–[34].TE Ma1 and Ma2 differ slightly by their regulons’ activity. TE Ma1 specifically activates the regulons *PRRX1*, *HES5*, *NFKB2*, *MIXL1*, *ATF7*, *IKZF1* and *XBP1* while TE Ma2 is more similar to TE Il1 and specifically activates the regulon *ZBTB33*, *GABPA*, *FOXO3*, *HAND1*, *NR6A1* and *USF2* (Figure 2D).

We then searched for enriched functions in TE populations, by performing a GSEA (Supplementary Table 3d). We observed that fatty acid anabolism, together with ovarian steroidogenesis, increased after TE differentiation (Figure 2E), whereas these functions of TE were not enriched at earlier stages of TE development.

### 3. Hypoblast specification and differentiation

To further characterise the HYPO cell population, we selected 15,335 cells from our dataset, corresponding to this population and performed dimensionality reductions followed by a new clustering (Figure 3A). This led to the identification of four HYPO subpopulations (Figure 3B). All of these subpopulations show a clear HYPO signature by expressing of *COL18A1*, *SOX17*, *GATA4* and *APOE* (Figure 3C and Supplementary Table 3a) [11], [35] but each one presents distinct features.

**Figure 3.**
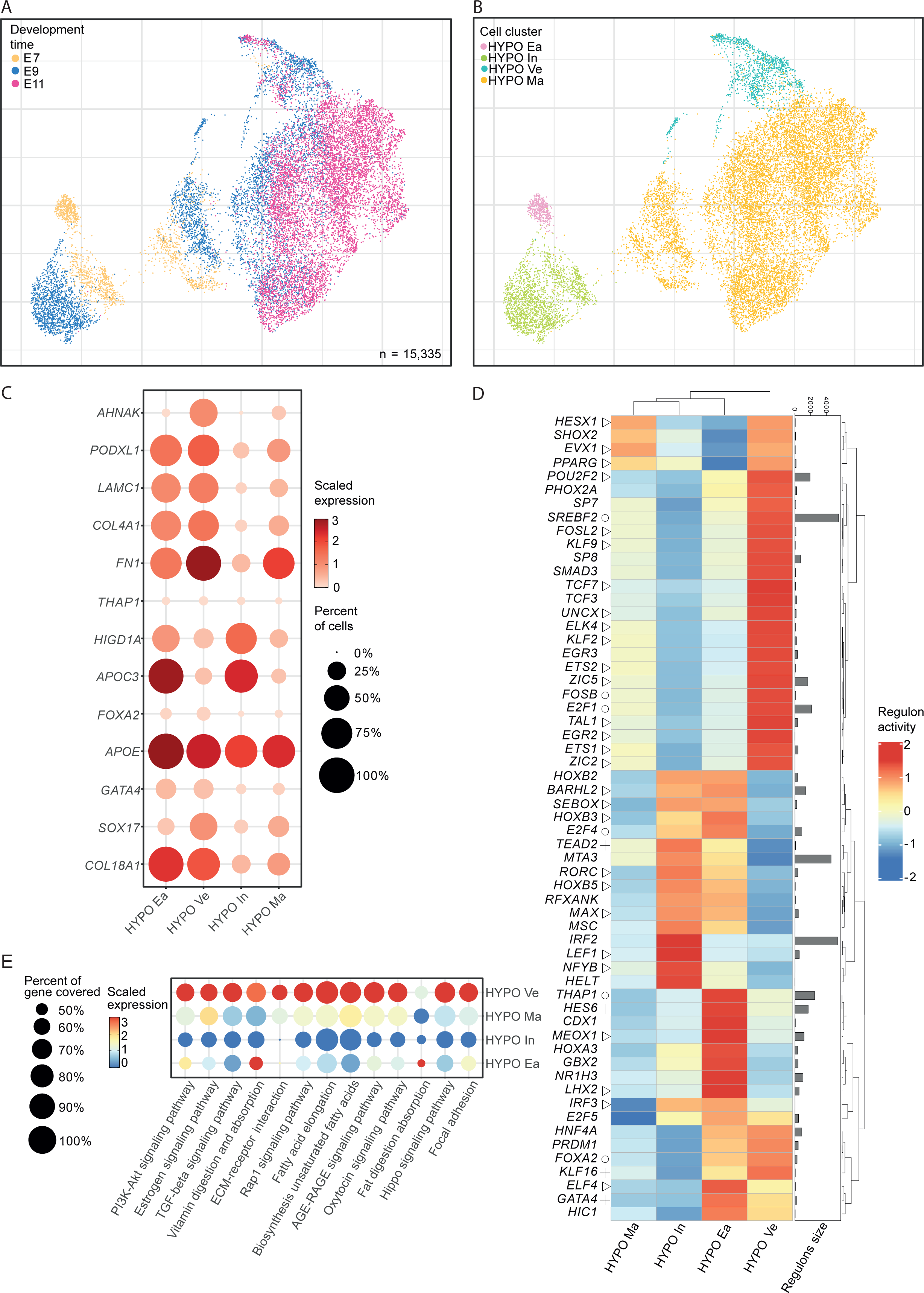
Identification and characterisation of different populations in the hypoblast. (A) Visualisation of selected HYPO cells coloured by developmental day via UMAP: E7 (yellow), E9 (blue) and E11 (pink). (B) Visualisation of HYPO populations coloured by cluster via UMAP: HYPO Ea (pink), HYPO In (green), HYPO Ma (yellow), HYPO Ve (blue). (C) Dot plot visualization of selected DEG genes, the circle size represents the percentage of cells within the cluster that express the gene. The red gradient represents the mean scaled expression of the genes within the cluster. (D) Heatmap showing scaled values of Regulon Activity Score for the 20 most specific regulons for each cluster, identified by Regulon Specificity Score (RSS). (△): common regulons with another pig study (+): common regulons with another human study; (Օ): common regulons within the three studies. Right rows (heatmap): histograms distribution of regulon size (number of genes regulated by the TF in the regulons). (E) Dot plot visualization of selected KEGG signalling pathways. The circle size represents the percentage of genes within all the genes in the pathways that express by the clusters. The red gradient represents the mean scaled expression of the pathways within the cluster, AGE-RAGE signalling pathway in diabetic complications have been truncated as AGE-RAGE signalling pathway.

These four clusters can be separated into two distinct groups according to developmental time and expression of specific genes (Figure 3A). The first group includes cells from E7 and E9 embryos and comprises early hypoblast (HYPO Ea, pink dots in Figure 3B) and intermediate hypoblast subpopulations (Hypo In, green dots in Figure 3B). It is characterised by the expression of *APOC3* and *HIGD1A* (Figure 3C and Supplementary Figure 5) and shares many active regulons (Figure 3D). The second group includes cells from E7 to E11 embryos and corresponds to mature hypoblast (HYPO Ma, yellow dots in Figure 3B) and visceral hypoblast (HYPO Ve, turquoise blue dots in Figure 3B). It is characterised by the expression of higher levels of fibronectin (*FN1)* and *COL4A1* (Supplementary Figure 5) and of genes associated with focal adhesion and ECM-receptor interaction (Figure 3E).

In the first group, HYPO In is characterised by an increased activity of the *IRF2*, *LEF1*, *NFYB* and *HELT* regulons, and some others common to the previously described TE Mt population, including regulons drivent by TFs of the *HOXB* gene family and *RORC.* HYPO Ea is characterised by an increased activity of regulons corresponding to TFs known to be early hypoblast marker genes (e.g., *FOXA2*, *THAP1* and *GATA4*) [36], but also regulons whose TFs are associated with the patterning of the anteroposterior axis (*CDX1*, *HES6*, *GBX2, MEOX1, HOXA3*) [37]–[39]. This first group could correspond to immature cell populations necessary for patterning the forming hypoblast.

The second group of cells consists of more differentiated hypoblast cells, with higher levels of genes associated with a mesenchymal phenotype (Figure 3E). These cells are also more cycling (supplementary Figure 3) with more than 60% and 70% of cells in G2/M and S phases for HYPO Ma and HYPO Ve, respectively, compared to less than 50% for the HYPO Ea. We also observed an increase in the expression level of genes associated with biological functions related to the biosynthesis of fatty acids and estrogen/oxytocin signalling pathways (Figure 3E). Within this group, HYPO Ma represents the majority of cells and should correspond to the parietal hypoblast, underlying the trophectoderm. The second HYPO population which was named Visceral Hypoblast (HYPO Ve) has DEG markers implicated in hypoblast differentiation (e.g., *GATA6*, *COL4A1*, *LAMC1*, *PODXL* and *AHNAK,* Supplementary Figure 5), which have been described in the derivation of extra-embryonic endoderm from pluripotent stem cells and in the regulation of hypoblast differentiation [35], [40]–[42]. This population presents a distinct regulon activity profile with HYPO-associated TFs (e.g., *FOSB*, *FOSL2*, *FOXA2*, *GATA*4) [43], but also with known targets of the Wnt signalling pathway (*TCF3*, *TCF7*), BMP/Nodal pathway (*SMAD3*) and with the activation of members of the *KLF*, *ETS*, *ZIC* and *EGR* family (*KLF9*, *KLF2*, *ETS1*, *ETS2*, *ZIC2*, *ZIC5*, *EGR2*, *EGR3*) known for their mitogenic and patterning activity (Figure 3D). We postulate that this population could be the one surrounding the embryonic disc and should correspond to the visceral endoderm described in mice. This hypothesis is supported by the expression of the Wnt signalling pathway inhibitors *DKK1* [44], which has been described in the mouse anterior visceral endoderm and *BMP2,* which has been shown to induce visceral endoderm differentation from XEN cells [45], [46] (Supplementary Figure 5).

### 4. Pluripotency states follow pig epiblast development

To further characterise the ICM and EPI cell populations, we selected 1,155 cells from our dataset corresponding to these populations and performed dimensionality reductions followed by a new clustering (Figure 4A). This revealed two distinct subpopulations (Figure 4B), which differ in terms of cell proliferation: more cycling cells were observed in the EPI compared to the ICM (84% vs. 60% in G2/M and S phases, Supplementary Figure 3).The first subpopulation is mainly composed of cells from the early blastocyst stage (E5) (Figure 4A) and corresponds to the ICM based on the expression profiles of genes associated with naive pluripotency (e.g., *ESRRB*, *KLF4*, *PDGFRA* and *STAT3* [11], [14], [15]) (Figure 4B-4C and Supplementary Table 3a). This population shares a high degree of transcriptional similarity with TE Ea, as indicated by their high similarity score (>0.97) (Supplementary Figure 7). This high similarity can also be explained by the ongoing active population segregation occurring between ICM and TE at this stage. In the regulon heatmap (Figure 4D), we identified early-stage regulons associated with ICM (e.g., *KLF17*, *STAT3*, *TCF7, NR2C2*) and reflecting the activity of known pathways (IL6/STAT3 and Wnt) associated with naive pluripotency in mammalian embryos [10], [11], [47]. We also found *ZNF148*, a TF known to suppress Notch signalling in induced pluripotent stem cells [48], [49] and *ZNF471* a TF described to affect stemness by down-regulating pluripotency markers (*NANOG*, *OCT4*, *SOX2*) [50] (Supplementary Figure 6).The second subpopulation, from E7 to E11, is distinct from the ICM and expresses known EPI population markers, associated with formative pluripotency genes (e.g., *NANOG*, *NODAL, DNMT3B*, *POU5F1*, *OTX2* and *SOX2,* Figure 4C). While this subpopulation appears to be quite stable along time, sharing a high degree of transcriptional similarity (Supplementary Figure 7), we observed a slight and graduate change in gene expression from E7 to E11, with an increase in the expression of *DNMT3*A, *LIN28A*, *NODAL* together with a decrease in the expression of *NANOG* and *STAT3* (Figure 4C and Supplementary Table 3a). This dynamic is also observed in terms of regulon activity with a decrease in activity for *NANOG*, *OTX2*, *GBX2*, *FOXD3*, *ETV1* and an increase in activity for *LHX1*, *LHX4*, *SMAD1*, *PROX1* and *PATZ1*. Our study also highlights novel regulons, whose functions regarding the biology of pluripotent stem cells, are not well understood. It includes *TFDP2, DBP, EN1, RFXANK*.

**Figure 4.**
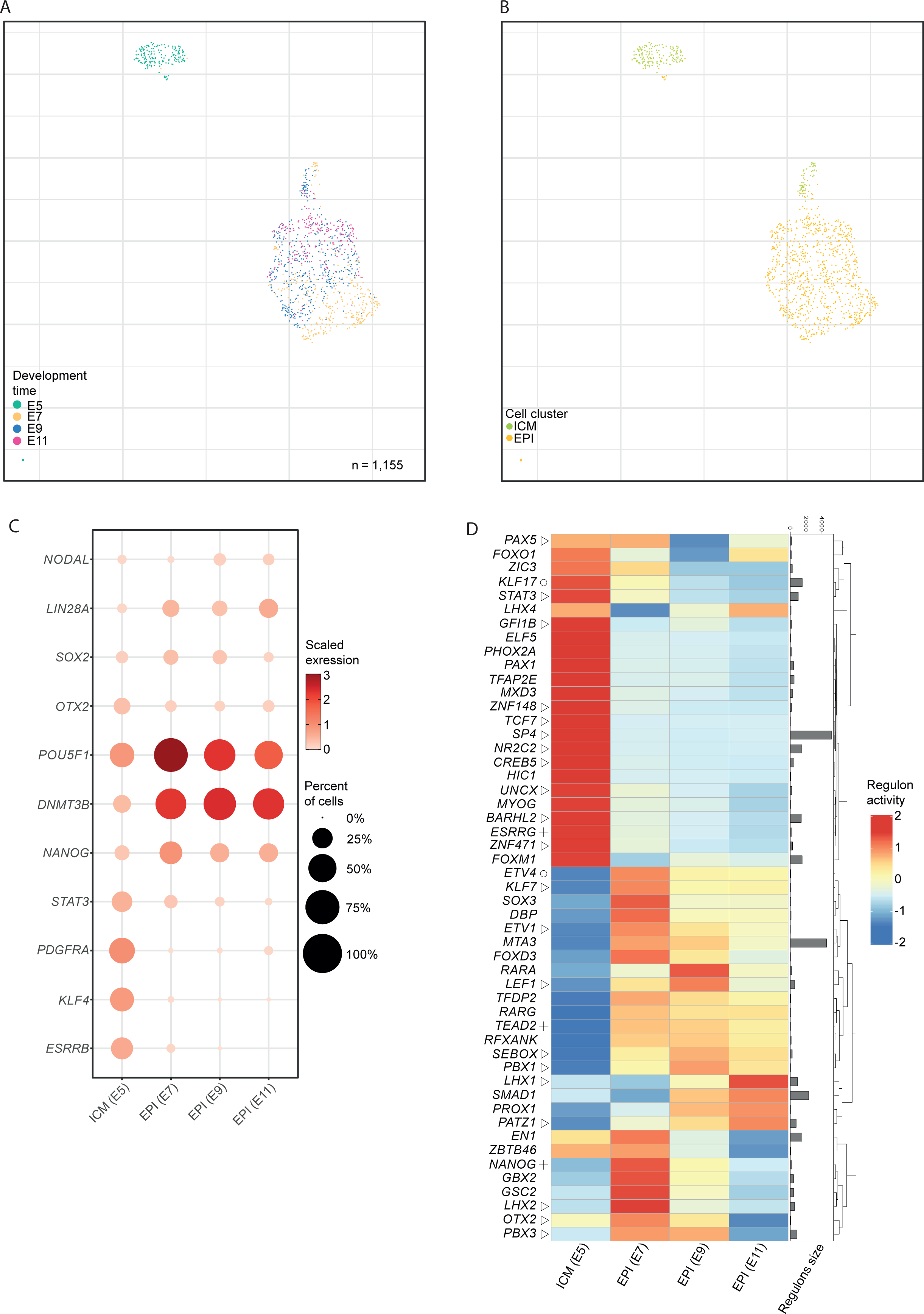
Characterisation of transcriptional change in epiblast populations. (A) Visualisation of selected ICM/EPI cells coloured by developmental day via UMAP: E5 (green), E7 (yellow), E9 (blue) and E11 (pink). (B) Identification of the two EPI populations coloured by cluster via UMAP: ICM (green), EPI (yellow). (C) Dot plot visualization of selected DEG genes. The circle size represents the percentage of cells within the cluster that express the gene. The red gradient represents the mean scaled expression of the genes within the cluster. (D) Heatmap showing scaled values of Regulon Activity Score for the 20 most specific regulons for each cluster, identified by Regulon Specificity Score (RSS). (△): common regulons with another pig study (+): common regulons with another human study; (Օ): common regulons within the three studies. Right rows (heatmap): histograms distribution of regulon size (number of genes regulated by the TF in the regulons. (E) Dot plot visualization of selected KEGG signalling pathways. The circle size represents the percentage of genes within all the genes in the pathways that express by the clusters. The red gradient represents the mean scaled expression of the pathways within the cluster.

### 5. Linking ligand-receptor interactions to regulon activation highlights potential functional interactions between the epiblast and the extra-embryonic populations

To link regulons to possible signalling pathways, we used the CellComm pipeline in order to create pathways by connecting ligand, receptor and TF based on their expression profiles. The network is based on known protein-protein interactions (Supplementary Table 5a) and the activity of each predicted pathway is scored using the average expression of TF, receptor and intermediates (Supplementary Table 5b). The main results are shown in Figure 5A. For the early blastocyst stage (E5), we found known pathways and players associated with naive pluripotency, acting either through a paracrine loop from the TE to the ICM (*ERBB2*, *FGFR4*, *PDGFRA*) or a paracrine/autocrine loop (TE to ICM or ICM to ICM) (*KIT*, *IL6ST*, *EGFR,* and *ITGB1*) (Figure 5). For the most active ones, these pathways converge to activate the downstream regulator *STAT3,* which is known to play a key role in the ICM and in naive pluripotent stem cells [10], [39]. We also observed the activation of *TCF7,* which is a direct target of canonical Wnt signalling and new pathways of interest, linking either *EGFR*, *IL6ST* or *ERBB2* to *ZNF148* or *EGFR* to *UBTF* (Figure 5A and 5B*). UBTF* has been described as a regulator of human ESC differentiation by regulating rRNA synthesis. Activin A treatment has also been shown to reduce the binding effects of *UBTF* [51] (Supplementary Figure 6).

At the subsequent E7 hatched blastocyst stage, the pathway activity profile shows no particular autocrine signalling for the EPI (Figure 5C), while signalling from TE and HYPO to EPI or TE occurs via *ITGB1*. TE also signals to EPI via *LRP2* and HYPO shows autocrine signalling occurring via *ITGA5,* particularly in HYPO Ve (Figure 5B).

At the subsequent E9 early ovoid blastocyst stage, paracrine and autocrine signalling by *ITGB1* and *ITGA5* are still predicted in HYPO and EPI but not between TE and EPI, where *CDH1* is mainly involved. We can also observe a weak signalling from EPI to TE via *PTPRF*, which is confirmed by the expression of *PTPRF* by TE Lr (Figure 5A and 5B). Interestingly *PTPRF* has been found in uterine extracellular vesicles of pregnant sows [52].

At the subsequent E11 late ovoid blastocyst stage, signalling from TE and HYPO converge on EPI, either through the activation of *ERBB3* or *ITGB1*. The two signalling pathways converge to activate *HNF4A*, *STAT3*, *HAND1*, and *SMAD1/3,* suggesting an importThe convergence of many signalling pathways towards EPI by ITGB1 supports the importance of the extracellular matrix and cellular contacts in transmitting the information necessary for the biology and survival of pluripotent cells. This may be a promising avenue of resant reactivation of signalling pathways crucial for early patterning of the embryonic disc and linked to the early steps of gastrulation.

earch to improve the derivation, stability and survival of porcine ESCs.

### 6. Changes in uterine fluids composition are associated with the transition between early and late blastocysts

To investigate potential functional interactions between the embryo and its surrounding uterine fluids (UFs), we sampled the UFs from the same sows used to produce the embryos. Uteri were flushed in order to recover the embryos and the uterine fluids. Uterine fluids were analysed by liquid chromatography-mass spectrometry (LC-MS/MS). A total of 1239 proteins were identified from which 277 were quantified in the 18 samples (Supplementary Table 6). At the quantitative expression level of the identified proteins, we found a clear discrimination of the proteins based on their LFQ intensities between early (5 days post-insemination (IA)) and late (9-11 days post-IA) UFs, clearly seen in the PCA (Figure 6A) as well as in the heatmap (Figure 6B). The early stage shows a protein intensity profile with functions associated with cell metabolism, such as those involved in glycolysis GAPDH, ENO1, AKR1A1, PKM, IDH1 (Figure 6B) [53]–[57] pyruvate mechanism LDHA/B (Figure 6B) [58] and proteins with pleiotropic functions such as proteins of 14-3-3 and YWHAQ/Z/E families, recently identified as key players during the maternal-to-zygotic transition in pigs (Figure 6B) [59].

**Figure 6.**
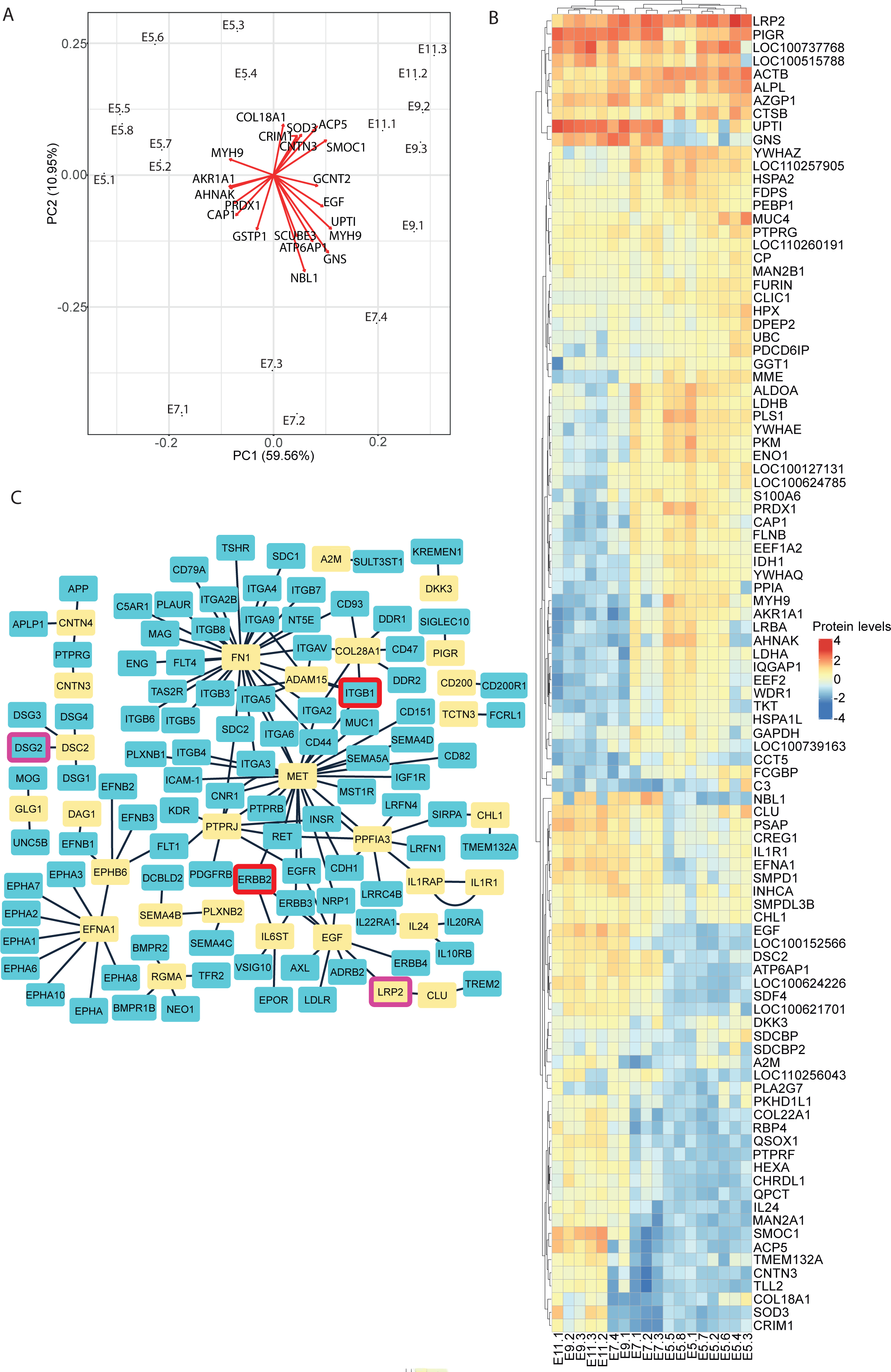
Change in the uterine fluids between E5 to E11 reflects potential crosstalk between the uterus and the embryo. (A) Principal Component Analysis (PCA) plot of pig uterine fluids. Each dot represents one uterine fluid sample. Sample IDs consist of embryonic day stages followed by the sample number. Red arrows represent prominently altered (negatively and positively) proteins for each of the two principal components (PC1 and PC2). (B) Heatmap showing scaled protein levels values of the 87 most prominently altered proteins. (C) Visualisation of possible interactions between uterine fluid proteins and receptor located on the TE or the EPI. Yellow boxes indicate proteins found in the uterine fluids, blue boxes indicate receptors, red circled boxes indicate receptor preferentially expressed in the EPI and purple circled box indicated receptors preferentially expressed in the TE.

In the later stages (E9-E11), classical markers of maternal-embryo recognition are detected, such as the interleukin complex with IL1RAP, IL24 and IL1R1 (Figure 6B) [60], [61]. We also detected receptors that were also identified in the TE, including UPTI, DAG1, PTPRF, DSC2 and LRP2 (Figure 5B, Supplementary Figure 4). Surprisingly, the JAK/STAT activator IL6ST was also identified (Supplementary Table 6). UPTI (uterine plasmin/trypsin inhibitor), DAG1 and LRP2 are also known to be expressed by endometrial cells [62]–[64], supporting the idea of a reciprocal loop of activation for similar signalling pathways between the endometrium and the conceptus to synchronise both for implantation.

By using ligand-receptor analysis, we then associated proteins detected in the UFs (in yellow) with previously identified receptors (in blue) expressed by EPI or TE cells from E9 to E11 embryos in the scRNAseq dataset (Figure 6C). In particular, we linked the expression of DSC2 in uterine fluid to the expression of DSG2 by TE cells. DSC2 has been found to interact with several desmoglein receptors and DSC2 is required for the formation of the blastocoel in bovine embryo [65]. We have also highlighted the importance of protein of the extra-cellular matrix like FN1, COL28A1, which can activate integrin signalling, like ITGB1 in EPI cells (Figure 6C and Figure 5A). We also observed the EPHA-Ephrin-A signalling pathway, with EPHA2 expressed by the pluripotent epiblast and EFNA1 expressed by the surrounding TE. EPHA-Ephrin-A plays an important role in the switch between pluripotency and differentiation in murine ESCs [66]. The presence of EFNA1 in uterine fluids at the onset of gastrulation also suggests that segregated EPH-EFN expression coordinates cell fate and early differentiation during early embryonic development. EGF is also enriched in UFs at E9-E11 stages and may interact with EPI and TE through different receptors (such as ERRB2 and LRP2, respectively), for self-renewal of pluripotent cells [67] and to prepare TE and endometrial cells for implantation [68]. We have also identified an interaction between A2M and SULT3ST1. A2M is expressed by the mouse endometrium and has an inhibitory effect on mouse blastocyst development [69]. For protein identified in endometrial development, we found changes between early stages with S100A6, CAP1 (Figure 6B) [70] and late stages with QSOX1, COCH, SMPD1 (Figure 6B) [71]–[73]. We also identified proteins annotated as negative regulators of BMP signalling pathway: NBL1, CHRDL1, SMOC1, CRIM1, COCH (Figure 6B) [74].

## Conclusions

Our study provides new insights into the formation of the first cell lineages of the early pig embryo from single-cell gene expression datasets from E5 to E11 embryos. In particular, our data reveal unsuspected dynamic evolution and heterogeneity within extraembryonic cell populations.

First, we confirmed the major difference between the early blastocyst at E5 and the later blastocysts from E7 onwards, both for ICM/EPI and early TE cells, whose transcriptional profiles are very different from the later lineages. This also suggests that the time window between E5 and E7 is crucial for the segregation of the three major lineages in the pig species and would deserve a more advanced and detailed analysis to understand the molecular mechanisms involved.

For the later blastocysts, whose biology differs from that of primates and rodents, we have highlighted previously unknown subpopulations, notably from E9 onwards, at the ovoid blastocyst stage. In the trophectoderm, we confirmed the existence of interleukin-1B secreting cells belonging to a very specific subpopulation, as well as known TE functions, such as lipid metabolism and catabolism. Above all, we discovered two previously unknown subpopulations of the TE. The first is characterised by the expression of LRP2, which could represent a subpopulation of TE progenitor or stem cells, and the second is characterised by the expression of numerous pro-apoptotic markers and disappears between E9 and E11. It could correspond to the cells of the Rauber’s layer.

Concerning the hypoblast, which we detected from E7 onwards, we observed two main groups of populations: some rather immature and present at E7 and E9 and others more mature and observed from E7 to E11. Among the latter, we find a population that could correspond to the visceral hypoblast, and another to the parietal hypoblast.

For the epiblast, we confirmed a quite stable pluripotent state from E7 to E11 with a graduate priming toward cell differentiation and gastrulation.

An original aspect of our study was to highlight regulatory modules (regulons) specific to each sub-population and potentially conserved between pigs and humans. Experimental validation will be required to confirm the relevance of such regulons in controlling the biology of pluripotent and extra-embryonic cells.

Finally, combined with the analysis of uterine fluid composition, we infer complex dialogs between the maternal environment and the cell lineages of the embryo and identified modules of cell signalling linked to TF regulated genetic modules.

## Methods

### 1. Production of pig embryos

All the embryos were produced at the INRAE experimental unit GenESI (Rouillé, France). All the metadata associated with the biological samples used in this study have been submitted to FAANG Data Portal (https://data.faang.org/home) and BioSamples (https://www.ebi.ac.uk/biosamples/) and are summarized in the supplementary Table 1.

Two distinct protocols were used for the production of pig embryos. A first batch of embryos (7 and 9 days after artificial insemination) was produced following superovulation (Supplementary information, Table 1). For each sow, estrous cycle was synchronized using Altrenogest (Regumate), a synthetic progestin during 18 days. The day after the end of the Altrenogest treatment, sows were superovulated using a first injection of gonadotrophin (1,200 UI PMSG) followed 72h later by an injection of 500 UI hCG. The day after, sows were then artificially inseminated and the insemination was repeated the following day. When the gestational time matched the embryonic stage to be sampled, surgery was performed as follow. The morning of the surgery, the sow was showered, and received an intramuscular injection of anesthetic (Ketamine, 10 mg/kg) and analgesic (Xylazine, 2 mg/kg) to calm her down. Then, the anesthesia mask was placed on her snout, the evaporator was put into operation for diffusion of the volatile anesthetic (Isoflurane 2%). Once the effectiveness of the general anesthesia was noted, a laparotomy was performed and the uterus was extracted from the abdominal cavity, clamped and the embryos were collected by retro-flushing of the uterine horns from the bottom of the horn (uterus) upwards (ovary) in 100 mL of physiological saline solution. The uterus was then replaced in the abdominal cavity, the wound was sutured and the animal was placed back into the recovery room. This procedure was authorized by the French ministry of higher education, research and innovation under the authorization number: Apafis#10376-20170623130698. The full protocol has been submitted on the FAANG Data Portal and is publicly accessible using the following link: https://api.faang.org/files/protocols/samples/INRAE_SOP_PLUS4PIGS_EMBRYOS_SAMPLING_PROTO1_20230131.pdf

A second batch of embryos was produced without superovulation (Supplementary information, Table 1). Sows were synchronized and inseminated as previously described. When the gestational time matched the embryonic stage to be sampled (5, 7, 9 and 11 days after artificial insemination), the sows were transported from the breeding unit (Rouillé, France) to the slaughterhouse (Nouzilly, France). They were stunned by electronarcosis and bled. The uterus was clamped and rapidly extracted from the abdominal cavity. Then, the embryos were collected into two tubes of 50 ml by retro-flushing of the uterine horns from the bottom of the horn (uterus) upwards (ovary) in 100 ml of physiological saline solution. The full protocol has been submitted on the FAANG Data Portal and is publicly accessible using the following link: https://api.faang.org/files/protocols/samples/INRAE_SOP_PLUS4PIGS_EMBRYOS_SAMPLING_PROTO2_20230131.pdf

### 2. Preparation of single-cell suspensions

Once recovered, embryos were staged, pooled and transported in Embryo holding Media (IMV Technologies) for E5 and E7 embryos or DMEM/F12 for E9 and E11 embryos to the molecular biology laboratory in a thermostatically controlled chamber at 38°C. At arrival, embryos were transferred to a 4-well dish and staged again under a stereomicroscope. When necessary, embryos of the same stage were then pooled together into a drop of DMEM/F12 or IMV Embryo holding media and processed for cell dissociation. The full protocol is publicly accessible on the FAANG Data Portal https://api.faang.org/files/protocols/samples/INRAE_SOP_PLUS4PIGS_EMBRYOS_DISSOCIATION_PROTO3_20230131.pdf. Briefly, the zona pellucida (ZP) of E5 embryos was removed by transferring embryos into drops of 0.5% of pronase for no more than 5 minutes. The ZP was removed by aspirating/repulse the embryo using a pipette tip. Then, the embryos were washed into drops of embryo holding media. All the embryos (E5 to E11) were next incubated in pre-warmed Accutase for 10 minutes then pre-warmed TrypLE for 10 minutes followed by mechanical dissociation by several rounds of aspiration/repulse a pipette tip.

### 3. Production of scRNAseq libraries and sequencing

Dissociated cells were washed in DMEM/F12, counted and resuspended in PBS-0.4 % BSA according to 10X Genomics protocol: Chromium Next GEM Single Cell 3’ Reagent Kits v3.1 CG000204 Rev D (for embryos sampled in 2021) or Chromium Single Cell 3’ Reagent Kits v2 CG00052 Rev F (for embryos sampled in 2017).

A BioAnalyzer (or FragmentAnalyzer) profile and a Qbit quantification were performed for each sample after cDNA amplification and after library amplification. The libraries were sequenced on an Illumina HiSeq 3000 (batch1) and BGI DNBSEQ-G400 (batch 2) to obtain 144M to 328M of raw reads per library. The full protocol has been submitted on the FAANG Data Portal and is publicly accessible using the following link: https://api.faang.org/files/protocols/experiments/INRAE_SOP_PLUS4PIGS_scRNASEQ_LIBRARIES_PROTO4_20230228.pdf

### 4. Production of an extended annotation to cover 3’ UTR from porcine transcripts

Raw sequencing files were mapped to the *Sus scrofa* genome (11.1) using the Nextflow pipeline TAGADA (github.com/FAANG/analysis-TAGADA). Gene positions were annotated as per Ensembl build 102 and genes were filtered based on their biotype annotation to only contains genes matching one of this category: protein-coding, long intergenic non-coding RNA, antisense, immunoglobulin or T-cell receptor. We then used the quant3p script (github.com/ctlab/quant3p) to extend genes in the 3’ exon for those where reads aligned past annotated genes. We used genome parameter set to 1,341,049,888 and bam file alignment. Some additional modifications to the annotation were performed including gene deletion, addition or positions change. These changes are summarized in the project’s GitHub repository (https://github.com/Goultard59/pig_embryo_scrnaseq/blob/master/1_generating_matrices/RE ADME_annotation_extension.md) and the gtf file is available there: https://github.com/Goultard59/pig_embryo_scrnaseq/blob/master/1_generating_matrices/Sus_scrofa.Sscrofa11.1.102_10_26.filtered.gtf. Then CellRanger (version 6.1.1) was executed on this annotation to produce count matrices.

### 5. Single-cell RNA sequencing analysis

Raw gene expressions were converted to a Seurat object using the Seurat R package (version 4.3.0) [75]. Cells were removed if they had more than 10,000 or fewer than 500-300 expressed genes, or if >25-10% of their UMIs were derived from the mitochondrial genome with adjustment between samples (Supplementary Table 2). After filtering, the gene expression matrices for each sample from the same stages were merged, normalized and scaled using Seurat’s function with the default settings. We then performed data integration using Harmony [76]. To reduce the dimensionality of this dataset, gene expression matrices were analysed by principal component analysis (PCA). The first 10-20 principal components were further used as input with adjustments between stages (Supplementary Table 2) for UMAP dimensionality reduction using the RunUMAP function with the selected number of principal components and default parameters. Clustering was conducted using the “FindClusters” function with stage-adjusted resolution parameters (Supplementary Table 2). Cell clusters in the resulting two-dimensional representation were annotated to known biological cell types according to curated known cell markers [11]. Cells from EPI, TE and HYPO were then selected and separately re-analysed for the dimensionality reduction (PCA and UMAP) and the clustering step (Supplementary Table 2).

### 6. Secondary analysis

Functional analysis (Gene Ontology, Human Phenotype Ontology, miRBase and Kyoto Encyclopedia of Genes and Genomes (KEGG) enrichment) was performed in order to obtain the most significant pathways for each cell population using the GSVA package [77] with parameters set to ssgea method and sz minimum of 1. A linear model was then applied to the output matrix, followed by empirical Bayes statistics for differential enrichment analysis between cell types (Supplementary Table S3c). Then, DEG analyses were performed in a pairwise fashion between the different stages (Supplementary Table S3b) and between cell populations (Supplementary Table S3a), using the FindMarkers function of the Seurat package, with a filter set at 0.05 for p-value and at 25% for the minimum percentage of cells expressing DEG in the given identity. Based on the resulting DEG for each cluster, an enrichment analysis was performed using the gost function from gprofiler2 package [78] with parameters set to *Sus scrofa* (Supplementary Table S3d).

### 7. Multiple Factor Analysis

Multiple Factor Analysis (MFA) represents an extension of PCA for the case where multiple quantitative data tables are to be simultaneously analyzed [79], [80]. As such, MFA is a dimension reduction method that decomposes the set of features from a given gene set into a lower dimension space. In particular, the MFA approach weights each table individually to ensure that tables with more features or those on a different scale do not dominate the analysis; all features within a given table are given the same weight. These weights are chosen such that the first eigenvalue of a PCA performed on each weighted table is equal to 1, ensuring that all tables play an equal role in the global multi-table analysis.

### 8. SCENIC analysis

In order to run SCENIC on the pig genome, a custom database for RcisTarget and GRNboost was built. Transcription factor lists were generated following the methods used for the AnimalTFDB 3.0 [81] with the pig Ensembl build 102. The motif to Transcription Factor annotation was adapted to pig by gene orthology using OrthoFinder [82]. The motif database was built using aertslab scripts (github.com/aertslab/create_cisTarget_databases) with the best transcript score for each gene of the genomes based on the 10kb upstream regions. The library of motifs used in this manuscript is comprised of 10,560 PWMs from several sources [83]. The SCENIC pipeline was run using the VSN pipeline with an aggregation of 10 runs [84] (github.com/vib-singlecell-nf/vsn-pipelines).

For each regulon, RSS was computed based on the specificity results [85] (Supplementary Table 4b). The 20 regulons with the highest RSS were selected for each cell type and each stage to produce the heatmap and the CellComm analyses. For heatmap production, the AUCell regulons activity were averaged for each cell type and scaled before plotting.

### 9. Meta-analysis comparisons

Processed scRNAseq expression matrix from Meistermann *et al.* (2021) datasets were downloaded from https://gitlab.univ-nantes.fr/E114424Z/meistermannbruneauetalprocessed using raw counts and annotations of cells. Raw sequencing reads (fastq files) from Zhi *et al.* (2021) [11] datasets were downloaded from Genome Sequence Archive (GSA) using accession number CRA003960. Reads were then split using sabre (https://github.com/najoshi/sabre) with the parameters pairedEnd mode and max mismatch of 2. We used the barcode list from the Supplementary information Table 1 of [11] as the input list of barcodes used for sabre. Reads were then trimmed using Trim Galore [86]. Finally, the reads were mapped using the same script used for the human dataset, available at (https://gitlab.univ-nantes.fr/E114424Z/SingleCell_Align) with an hisat index based on 11.1 pig reference genome, a pig annotation reference ENSEMBL 104 and pairedEnd mode.

For SCENIC analysis, the previous pipeline was applied to Zhi *et al*. datasets (Supplementary Figure 9). For Meistermann *et al.,* we applied the previous pipeline with human adaptation as the v9 Motif2TF annotations and the feather file-based hg38 with TSS+/-10kbp was used (Supplementary Figure 8). Then a comparison between the transcription factor identified in those two and ours was performed (Supplementary Figure 10).

### 10. Ligand Receptor analysis

Ligand to transcription factor pathways was inferred using CellComm from FUSCA [87]. Transcription factors were retrieved from our previous SCENIC analysis. Ligands and receptors were identified with LIANA packages [88] where gene names were converted from humans to pigs with OmniPath [89]. The package was run to use sca, natmi, logfc, connectome, call_italk and call_connectome methods. Inference between all population at all stages was made with default parameters following CellCom tutorials. Protein-protein interaction was obtained from OmniPath with the conversion from human to pigs (Supplementary Table 5a and 5b). After interactions inference, the top 20 ranked pathways by possible autocrine or paracrine interactions were kept for visualization.

In order to produce circos plot visualization of interactions between the different populations, we considered the ligand to receptor activity score (Supplementary Table 5a) corresponding to the matching criteria. Ligand and receptor were then assigned to a population for which sender cell types show an expression that is higher than the average expression plus the standard deviation. Unassigned ligands and receptors were subsequently discarded. Then, circos plots were produced using the sum of the mean ligand and receptor expressions.

### 11. LC-MS/MS analysis of uterine fluids

Uterine fluids were recovered together with porcine embryos, by retro-flushing of the uterine horns from the bottom of the horn (uterus) upwards (ovary) in 100 ml of physiological saline solution. The embryos were removed and the resulting solution was then centrifugated and the supernatant filtered through a 70µm cell strainer and stored at −80°C before being processed.

Samples were concentrated using centrifugal filter devices (Amicon-4 10K, Merck) at 4,000 g for 25 min. Aliquots of concentrate were brought to a concentration of 1 M urea / 50 mM ammonium bicarbonate. Sample were then reduced in 5 mM DTE for 30 min at 37° C and subsequently alkylated in 15 mM iodoacetamide at room temperature in the dark for 30 min. Samples were digested with 70 ng of modified porcine trypsin (Promega) at 37° C overnight. Peptides were dried using a vacuum concentrator and resuspended in 0.1 % formic acid. Peptide samples were analyzed using a system consisting of an Ultimate 3000 nano-LC system online coupled to a Q Exactive HF-X mass spectrometer (Thermo Fisher Scientific). Peptides were injected on a PepMap 100 C18 trap column (100 µm × 2 cm, 5 µM particles, Thermo Fisher Scientific) and separated with an EASY-Spray analytical column (PepMap RSLC C18, 75 µm × 50 cm, 2 µm particles, Thermo Fisher Scientific). Chromatography was performed at flow rate 250 nl/min with 0.1% formic acid as solvent A and 0.1% formic acid as solvent B. The chromatographic method consisted of i) a 10 min equilibration step with 3 % solvent B, ii) a 90 min gradient from 6% to 20% solvent B, iii) a 10 min gradient from 20% to 40% solvent B and iv) a 10 min final elution step at 85 % solvent B. MS spectra were acquired using a top-15 data-dependent acquisition method on a Q Exactive HF-X mass spectrometer.

Protein identification and quantification was performed with MaxQuant (v.1.6.1.0) and the NCBI RefSeq *Sus scrofa* database. For the database search, the following parameters were used: enzyme: Trypsin/P; missed cleavages ≤ 2; 4.5 ppm mass tolerance for precursor main search; 20 ppm mass tolerance MS/MS search; carbamidomethylation of cysteine as fixed modification and acetyl (Protein N-terminus) as well as oxidized methionine as variable modifications. Label-free quantification was used.

### 12. Proteomics data analysis

The output table proteinGroups.txt from MaxQuant was loaded into Perseus [90] v.1.6.7 for downstream analyses. Data were filtered to remove contaminants, reverse peptides that match a decoy database and proteins identified only by modified peptides. The matrix was normalised by log^2^, and the samples were categorized in three categories early (E5), intermediate (E7) and late (E10). Proteins were kept if there were present in 70% of at least one category. Finally, missing values were replaced by a normal distribution (Supplementary information, Table S6). To assign protein names, we used the most identified protein IDs (Majority Protein IDs from MaxQuant output), then converted the RefSeq Protein Accession to gene symbol and ENSEMBL gene pig using gprofiler. The remaining ambiguous proteins were LOC100521789, assigned to AMY2, LOC110259139, assigned to PCDHA11, and ARF3, assigned to ARF1. PCA were performed using the R function autoplot from the package ggfortify [91]. Differential expression tests between early and late categories were produced using Perseus Volcano plot function with parameters t-test, both sides, 250 randomizations, an FDR threshold of 0.05 and a S0 of 0.1. Interaction between uterine fluids and receptor from TE and EPI were produced using LIANA: single cell data from EPI and TE at stages E9 and E11 were pulled together with 200 « artificial cells » produced from protein differentially enriched in the late categories. All cells were processed on LIANA using OmniPath ressources and orthologue genes converted using gprofiler and methods connectome, logfc, natmi, sca, cellphonedb, cytotalk, call_squidpy, call_cellchat, call_connectome, call_italk. Results were then filtered to keep only interactions between uterine fluids as ligand and TE and EPI cell populations as receptors. Resulting interactions were visualized anZhi et ald plotted using Cytoscape [92]. Receptor were assigned to a cell population based on their appearance on the differential expression testing between EPI and TE.

### 13. Data availability

All raw sequencing data and associated metadata are available in FAANG Data portal (https://data.faang.org/home) and ENA under accession number PRJEB60517. The code used for the analysis is available at (github.com/Goultard59/pig_embryo_scrnaseq/). The mass spectrometry proteomics data have been deposited to the ProteomeXchange Consortium via the PRIDE[93] partner repository with the dataset identifier PXD042421.

## Acknowledgments

We are grateful to the genotoul bioinformatics platform Toulouse Occitanie (Bioinfo Genotoul, https://doi.org/10.15454/1.5572369328961167E12) for providing help, computing and storage resources. We are grateful to people from the INRAE experimental farm (https://doi.org/10.15454/1.5572415481185847E12) who took care of the animals. We are grateful to Sylvain Bourgeais and the people from the slaughterhouse of the Unité Expérimentale de Physiologie Animale de l’Orfrasière INRAE (https://www6.val-de-loire.inrae.fr/uepao). We thank Marie-Jose Mercat from the IFIP, Institut du porc for her helps and funding. We also thanks Sarah Djebali, Nathalie Beaujean, Nathalie Vialaneix, Camille Berthelot and Laurent David for sharing data, codes and pipelines and for their helpful suggestions. We thank M. Kösters at LMU for excellent technical assistance. We thank people from our respective laboratories for helpful comments and discussion.

## Contributions

J.S., Y.B., P.M., F.M., T.F., J.A. and H.A. performed experiments. A.D., C.K., J.S., D.L., T.F., S.F., J.A. and H.A. analysed the data and prepared figures. A.G.T., S.F., B.P. provided reagents, technical expertise and conceptual inputs. A.D., S.F., J.A. and H.A. wrote the paper.

## Fundings

This study was funded by the ANR PluS4PiGs (ANR-19-CE20-0019). Adrien Dufour is partly funded by the DIM-1HEALTH from Région Île-de-France, the Animal Genetics division of INRAE and the IFIP, Institut du porc.

---

**Supplementary Figure 1:**
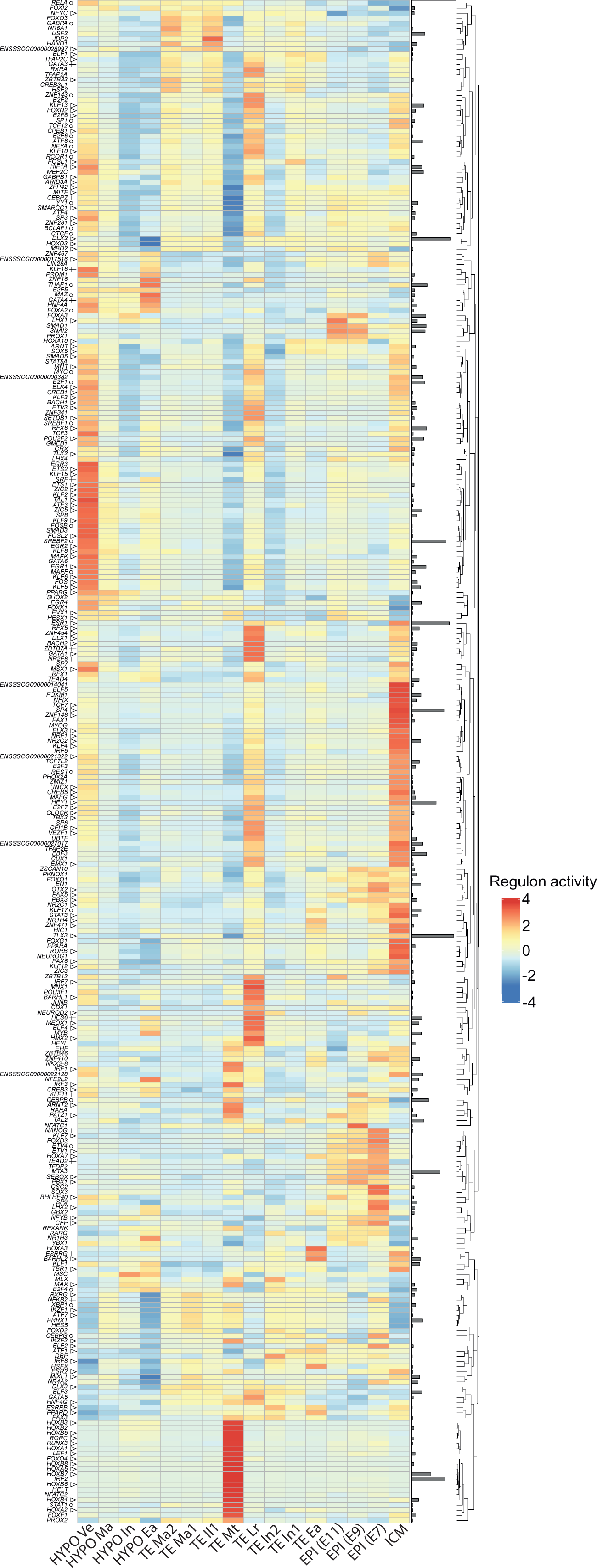
Heatmap of regulons in the different clusters. Heatmap showing scaled values of Regulon Activity Score for all the identified regulons, for each cluster. Regulons for transcription factors found in our meta-analysis of two other studies: triangle (△) for common regulons with another pig study, plus sign (+) for common regulons with another human study and circle (Օ) common regulons within the three studies. Right rows (heatmap): histograms distribution of regulons size (number of genes regulated by the TF in the regulons).

**Supplementary Figure 2:**
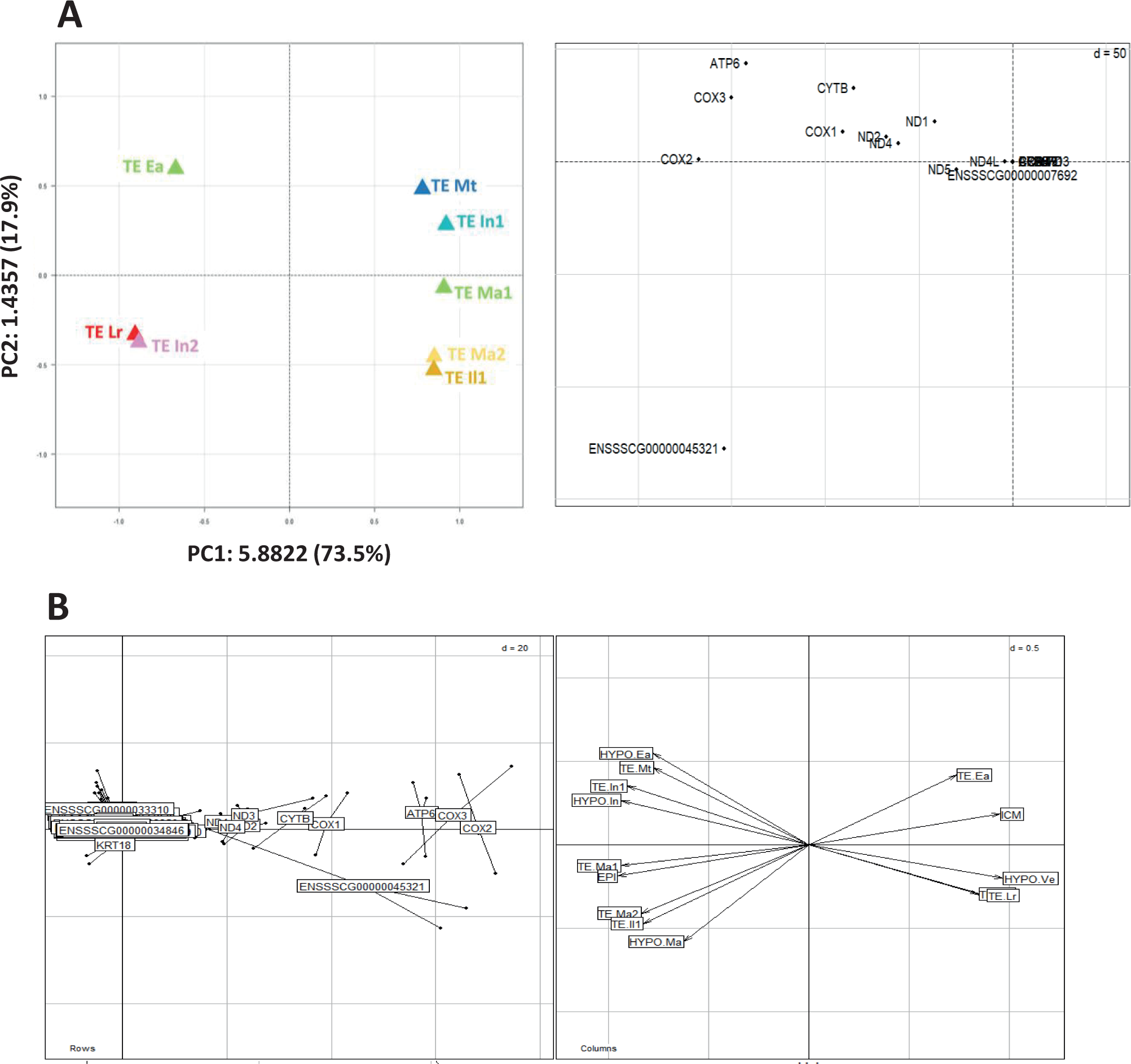
Principal Component analysis (PCA) and Multiple Factorial analysis (MFA) of genes and cell populations. **A: PCA of TE cell populations and genes.** The first axis (75.3% of the explained variance) separates populations regarding their metabolic phenotypes, as shown by the main contributions of genes linked to mitochondrial respiration (*ATP6*, *COX1/2/3*, *CYTB*, *ND1/2/3/4*). **B: MFA of genes and cell populations**. A clear dichotomy is seen within all the cell populations with common gene markers linked to mitochondrial respiration (*ATP6*, *COX1/2/3*, *CYTB*, *ND1/2/3/4*).

**Supplementary Figure 3:**
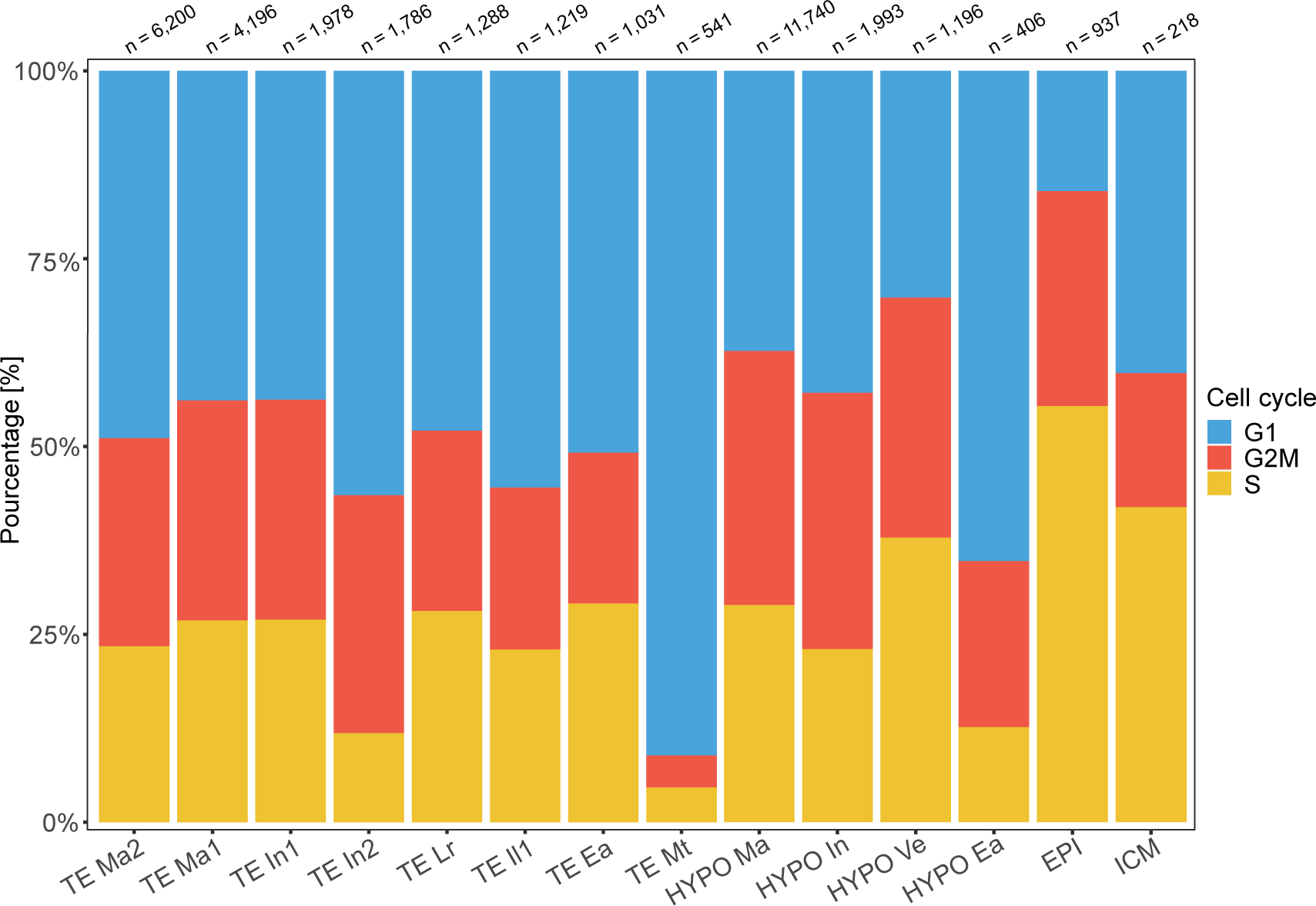
Cell cycle of the different cell clusters. Bar plot visualisation of the percentage of cells in each cell cycle phase: G1 (blue), G2M (red), S (yellow). Cell cycle phases were assigned to each cell using the Seurat CellCycleScoring function. Number of cells in each cluster is shown at the top.

**Supplementary Figure 4:**
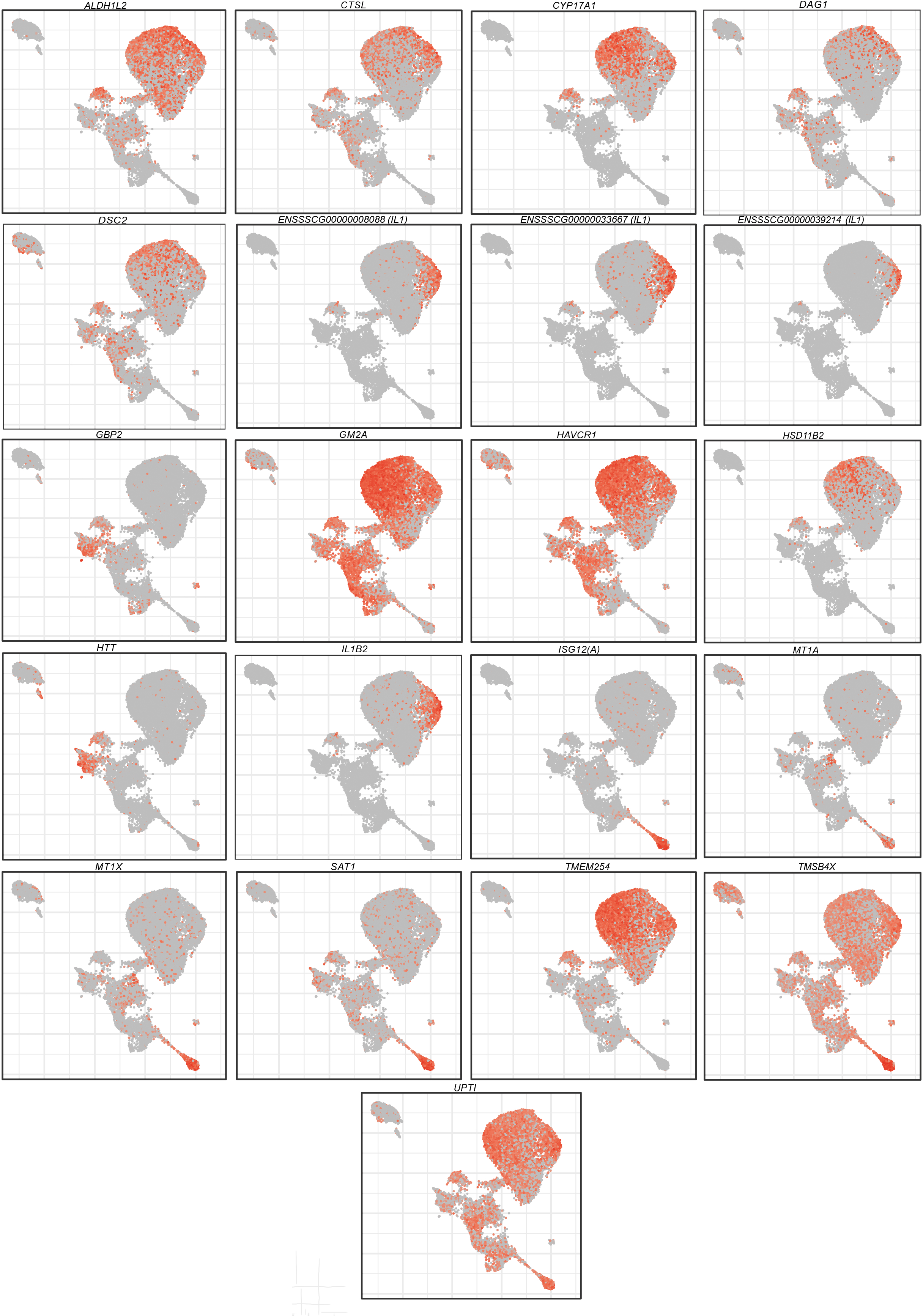
UMAP visualisation of selected genes in TE. UMAP visualisation of *ALDH1L2*, *CTSL, CYP17A1*, *DAG1*, *DSC2*, *ENSSSCG00000008088*, *ENSSSCG00000033667, ENSSSCG00000039214*, *GBP2*, *GM2A*, *HAVCR1*, *HSD11B2*, *HTT*, *IL1B2*, *ISG12(A)*, *MT1A*, *MT1X, SAT1*, *TMEM254*, *TMSB4X*, *UPTI* in TE cells.

**Supplementary Figure 5:**
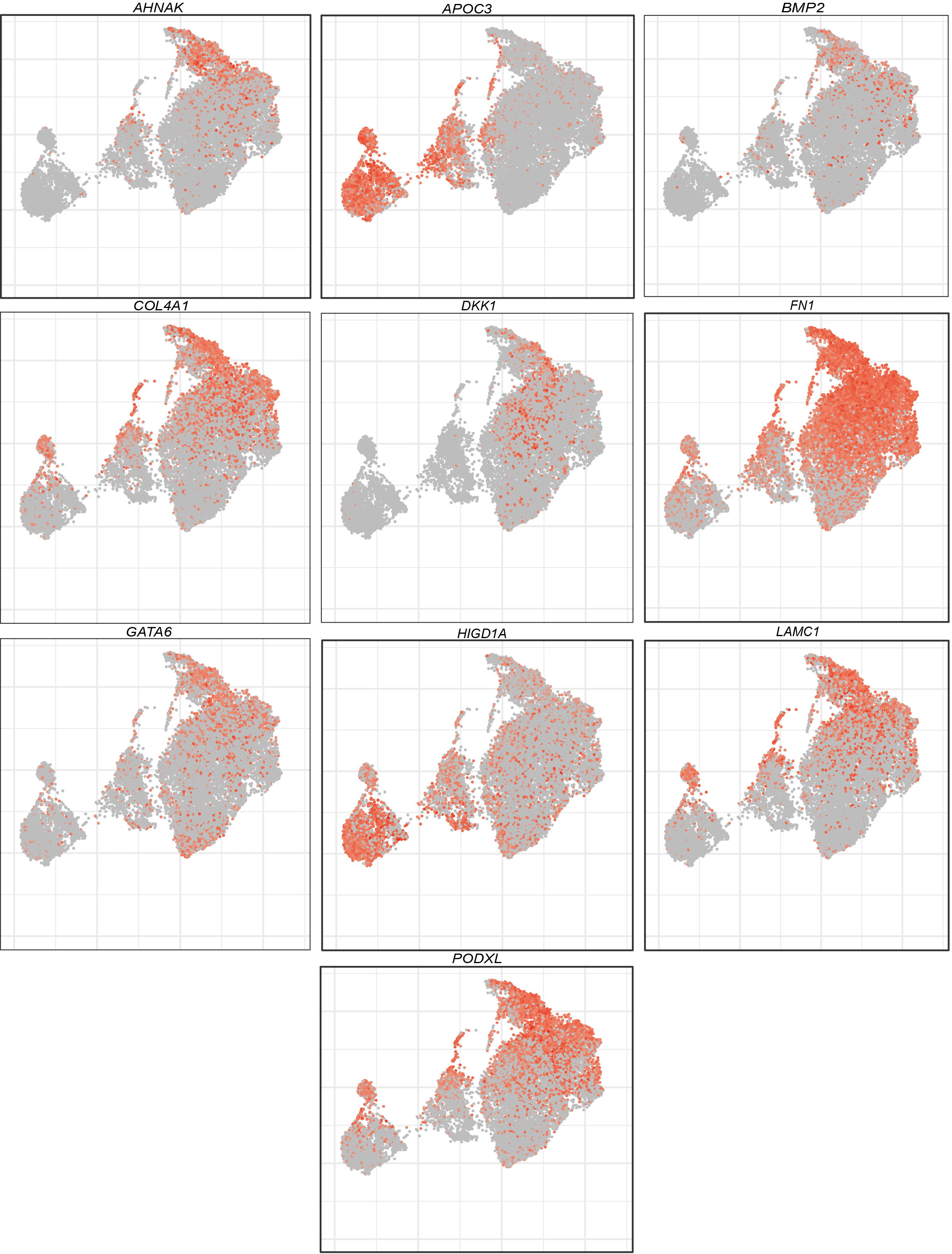
UMAP visualisation of selected genes in HYPO. UMAP visualisation of *AHNAK*, *APOC3*, *BMP2*, *COL4A1*, *DKK1*, *FN1*, *GATA6, HIGD1A*, *LAMC1, PODXL* in HYPO cells.

**Supplementary Figure 6:**
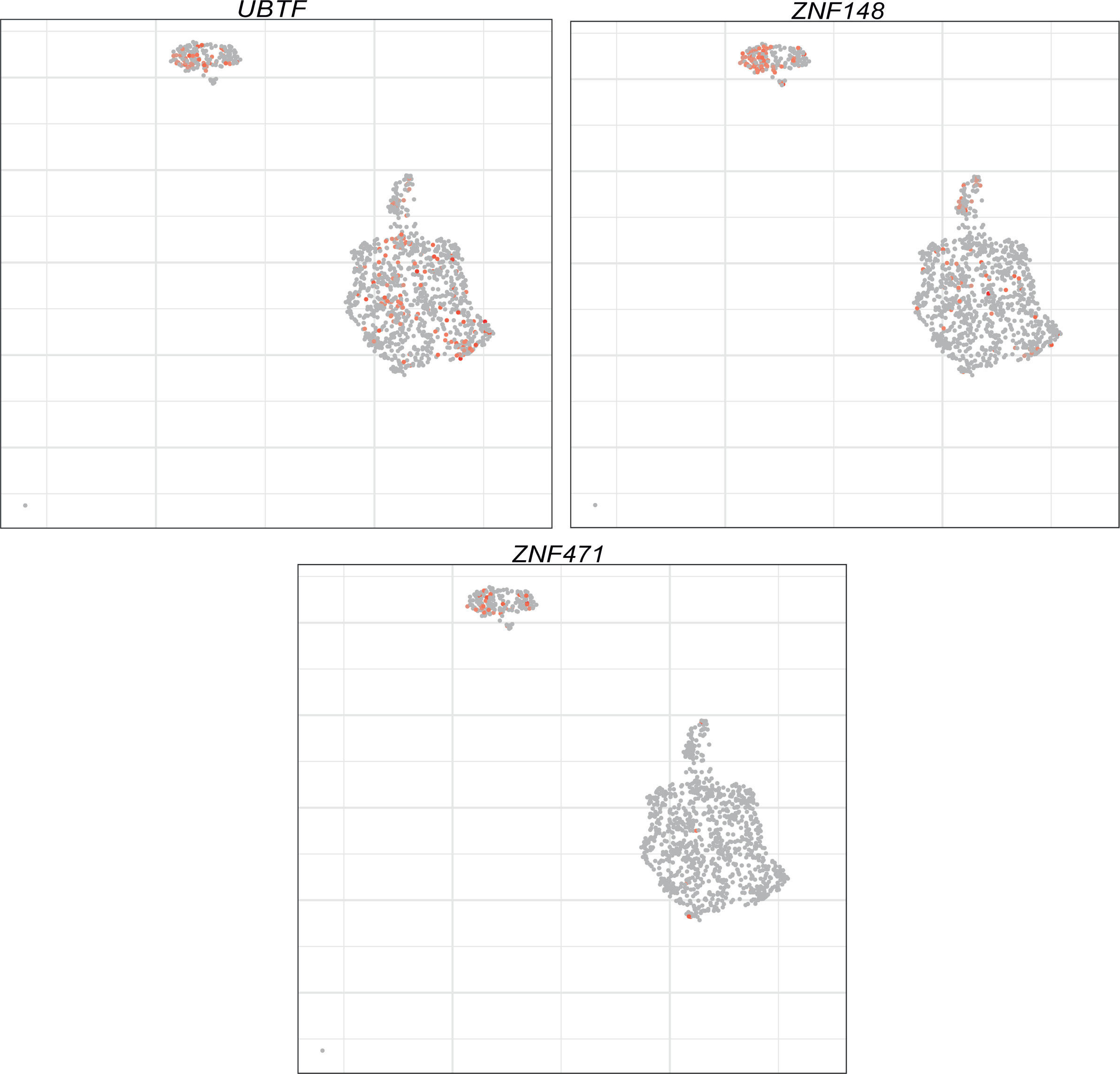
UMAP visualisation of selected genes in ICM/EPI. UMAP visualisation of *UBTF*, *ZNF148*, *ZNF471* in ICM/EPI cells.

**Supplementary Figure 7:**
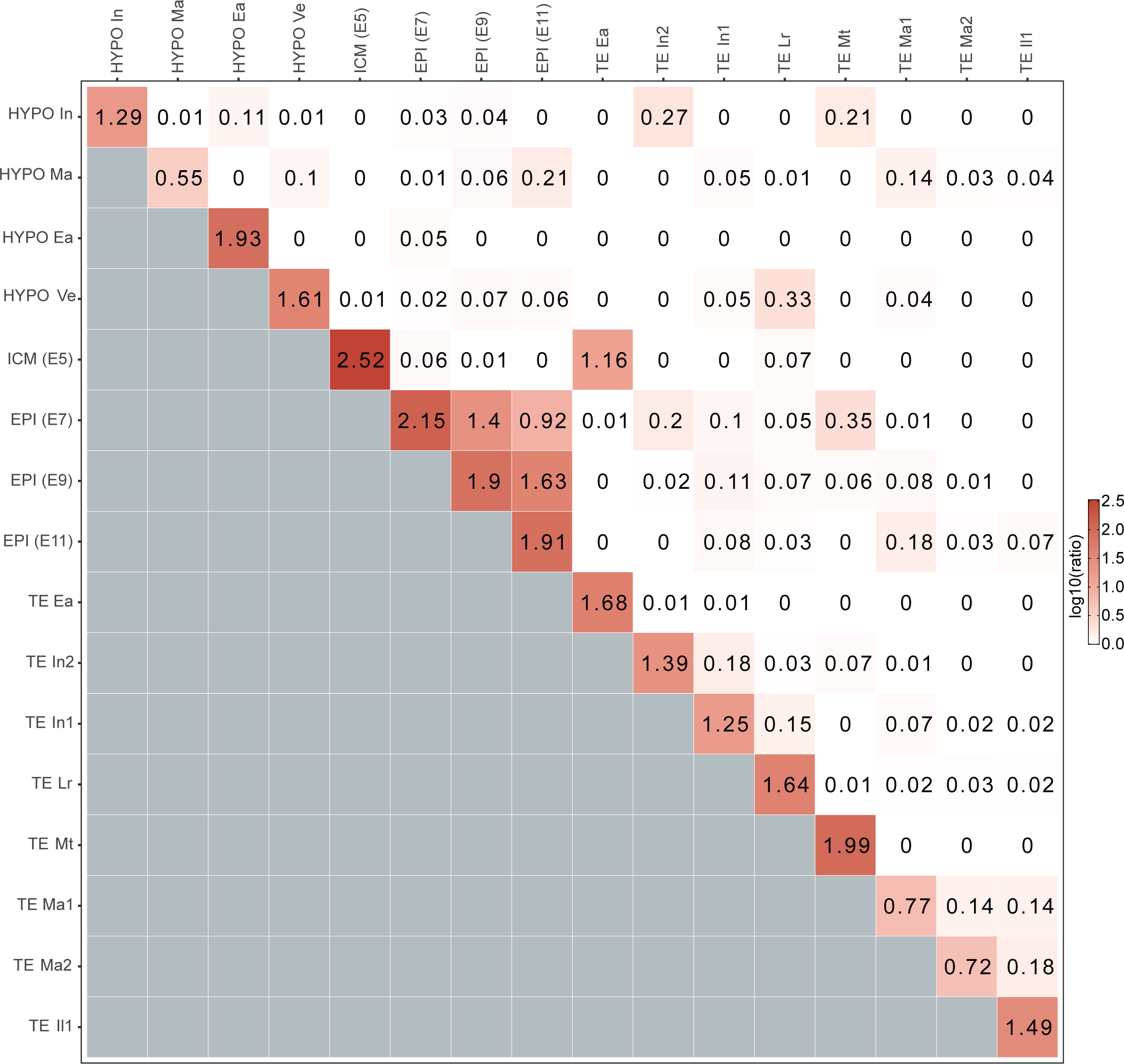
Matrix of cluster similarity. The correlation matrix was generated using the first 25 PCA on the total processed stage matrix. Then a nearest-neighbour graph was built using scran buildSNNGraph function (Lun ATL 2016), the matrix was produced using the pairwiseModularity function from the bluster package. The result has been converted by: log_10_(*x* + 1).

**Supplementary Figure 8:**
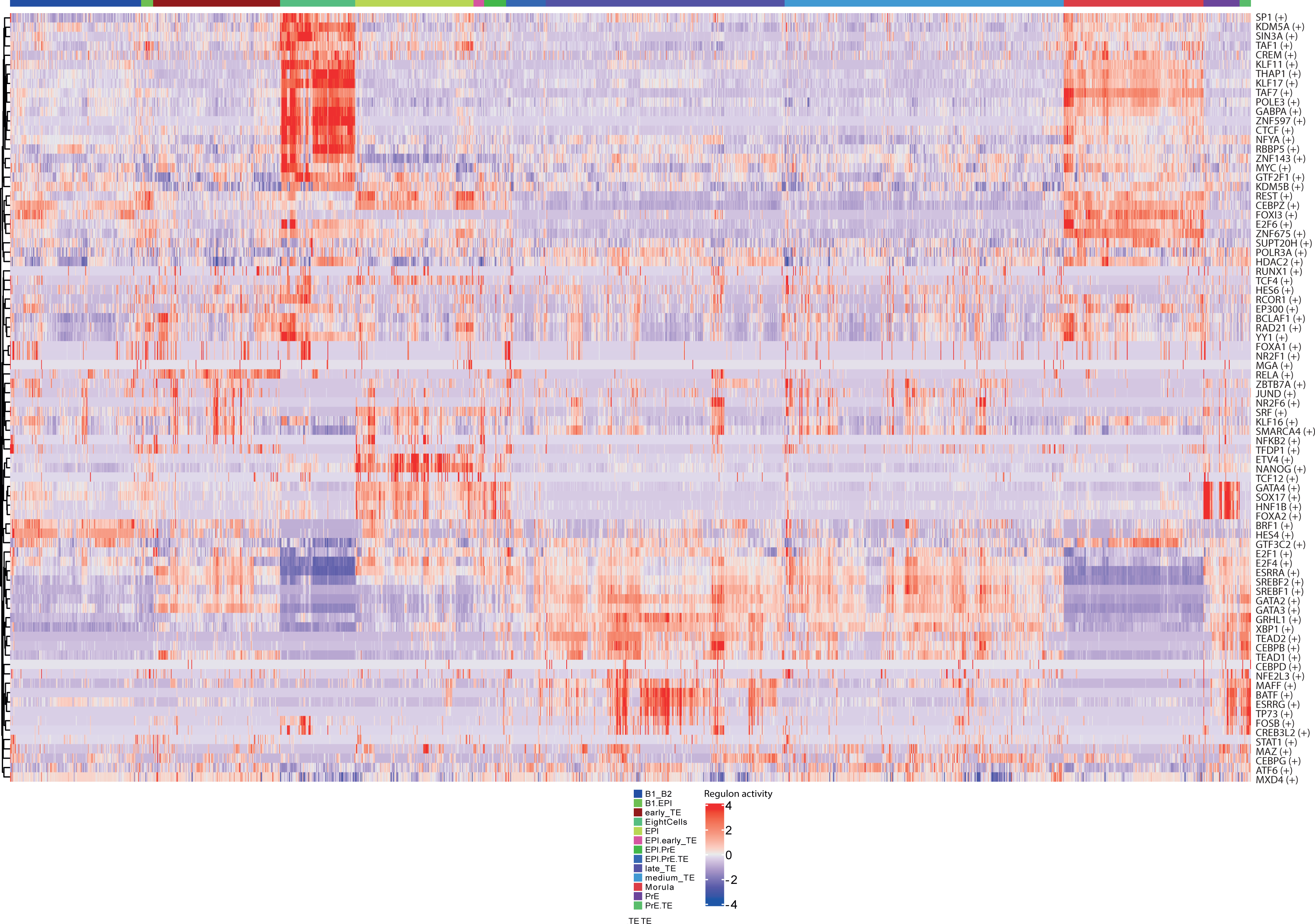
Heatmap of regulons in the different clusters from Meistermann et al. (2021) dataset. Heatmap showing scaled values of Regulon Activity Score for all the identified regulons, for each cluster described in the Meistermann et al. publication.

**Supplementary Figure 9:**
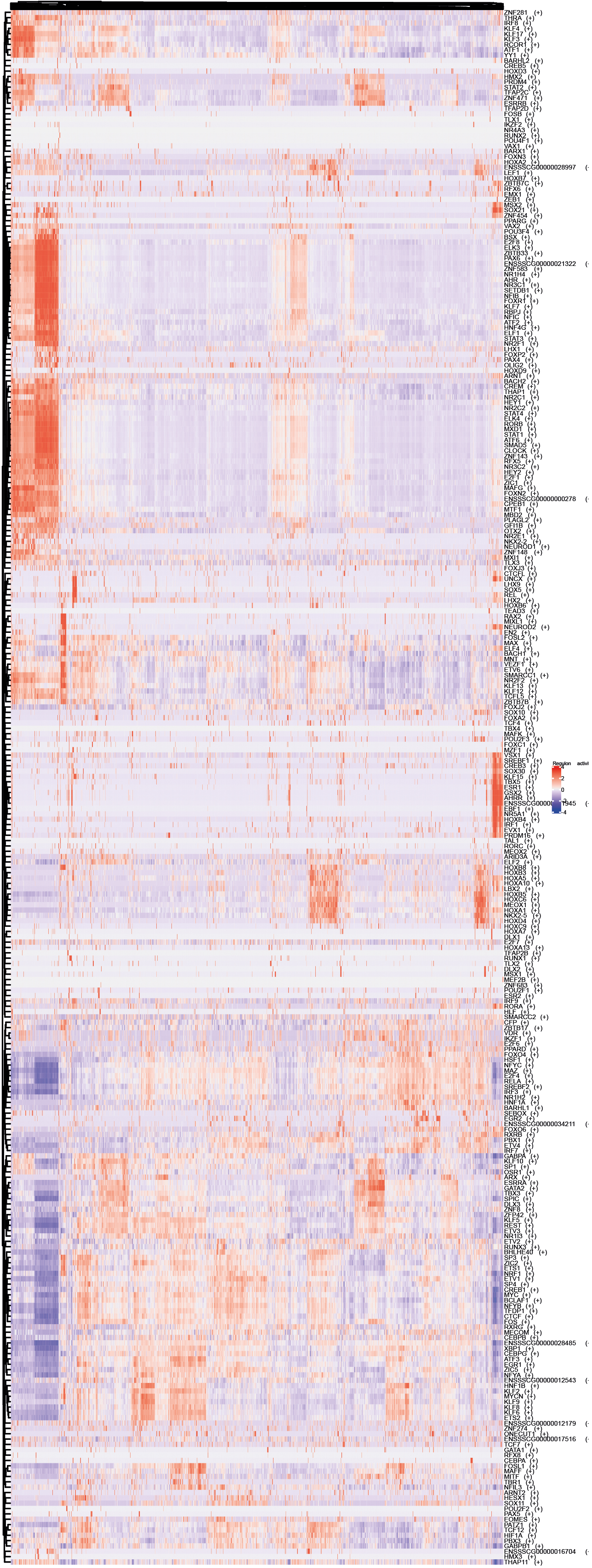
Heatmap of regulons in the different clusters from Zhi et al. (2021) dataset. Heatmap showing scaled values of Regulon Activity Score for all the identified regulons, for each cluster described in the Zhi et al. publication.

**Supplementary Figure 10:**
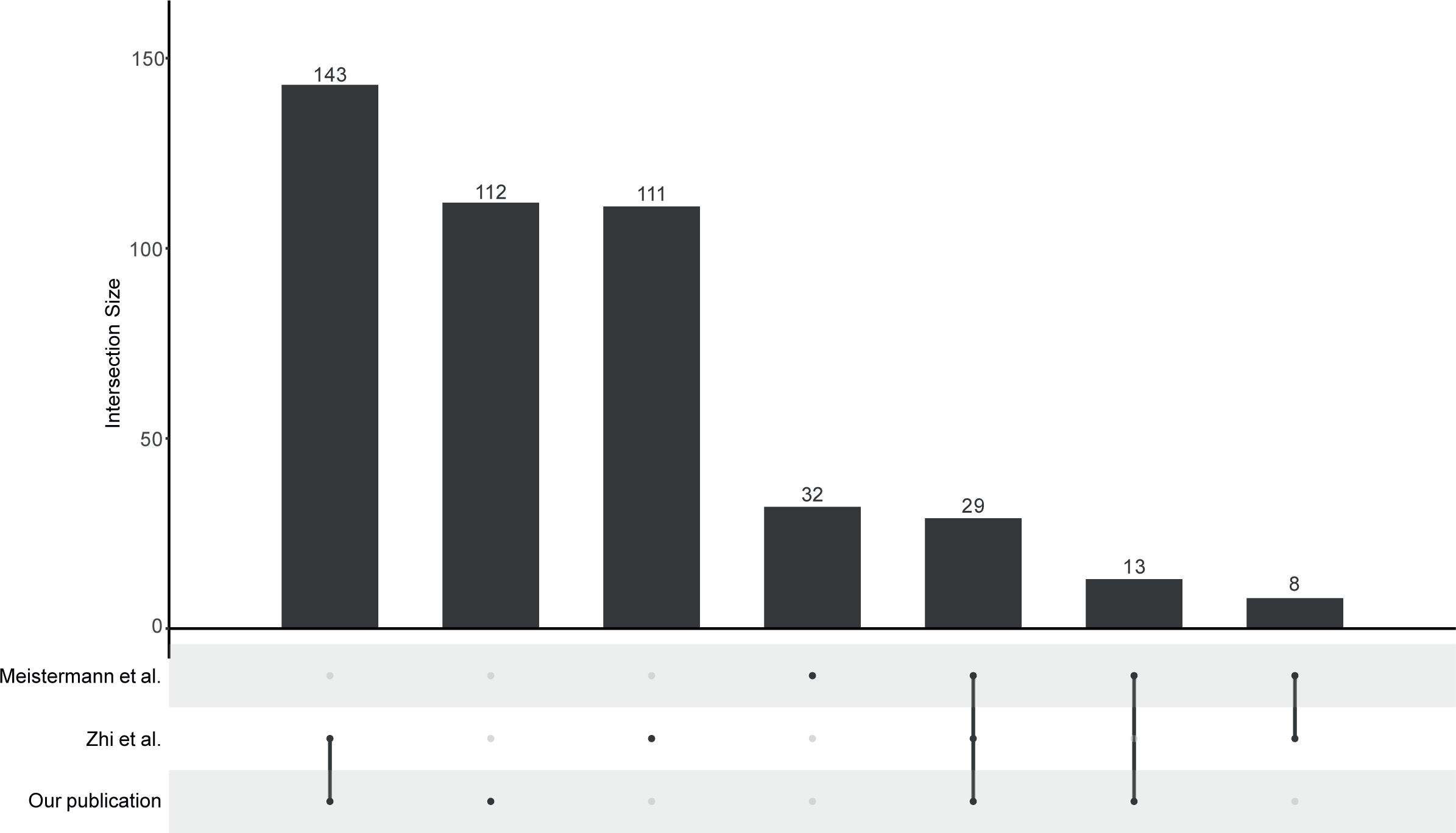
Upset plot of the comparaison of the identified transcription factor identified with SCENIC. Upset plot of the regulons identified by SCENIC between our study and the publications of Meistermann et al. (2021) and Zhi et al. (2021).

## Notes

### Competing Interest Statement

The authors have declared no competing interest.

