## Supplementary Table 1 for "Cell specification and functional interactions in the pig blastocyst inferred from single cell transcriptomics"

| Sow (ID & breed) | Date of sampling | BioSample ID sow | Boar (ID & breed) | Protocol for embryo production | Uterine fluid (sample name) | Stage | Number of embryos | BioSamples’ name (embryos & single cell specimen)  (prefix: INRAE_Plus4PigS_) | BioSamples’ IDs (embryos & single cell specimen) | Stage (description) | Library name :  (prefix : INRAE_Plus4PigS_ |
| --- | --- | --- | --- | --- | --- | --- | --- | --- | --- | --- | --- |
| 324  Large White | 30/08/2017 | SAMEA112465579 | 503240  Pietrain | superovulation | No | E7 | 17 | embryo_D7_1  embryo_single_cell_D7_1 | SAMEA112465608  SAMEA112465652 | hatched blastocyst | scRNAseq_E7_1 |
| 710237  crossed | 14/11/2017 | SAMEA112465584 | 501208  Pietrain | superovulation | No | E7 | 6 | embryo_D7_5  embryo_single_cell_D7_2 | SAMEA112465615  SAMEA112465653 | hatched blastocyst | scRNAseq_E7_2 |
| 710547  crossed | 14/11/2017 | SAMEA112465585 | 501208  Pietrain | superovulation | No | E7 | 10 | embryo_D7_6  embryo_single_cell_D7_3 | SAMEA112465616  SAMEA112465654 | hatched blastocyst | scRNAseq_E7_2 |
| 640  Large White | 22/08/2018 | SAMEA112465587 | 808055  Duroc | superovulation | No | E9 | 3  1 | embryo_D9_5  embryo_single_cell_D9_1  embryo_D9_4  embryo_single_cell_D9_2 | SAMEA112465619  SAMEA112465655  SAMEA112465618  SAMEA112465656 | ovoid blastocyst D9 10mm | scRNAseq_E9_1  scRNAseq_E9_2 |
| 2292  Large White | 27/01/2021 | SAMEA112465588 | 2000966  crossed | normal breeding procedure | Yes  2292_E5-4 | E5 | 10 | embryo_D5_1  embryo_single_cell_D5_5 | SAMEA112465620  SAMEA112465661 | early blastocyst (with blastocoel) | scRNAseq_E5_2 |
| 2248  Large White | 27/01/2021 | SAMEA112465589 | 2000966  crossed | normal breeding procedure | Yes  2248_E5-8 | E5 | 19 | embryo_D5_2  embryo_single_cell_D5_1 | SAMEA112465621  SAMEA112465657 | early blastocyst (compact) | scRNAseq_E5_1 |
| 2195  Large White | 27/01/2021 | SAMEA112465590 | 2000966  crossed | normal breeding procedure | Yes  2195_E5-7 | E5 | 18 | embryo_D5_3  embryo_single_cell_D5_2 | SAMEA112465622  SAMEA112465658 | early blastocyst (compact) | scRNAseq_E5_1 |
| 2391  Large White | 27/01/2021 | SAMEA112465591 | 2000966  crossed | normal breeding procedure | No | E5 | 10 | embryo_D5_4  embryo_single_cell_D5_6 | SAMEA112465623  SAMEA112465662 | early blastocyst (with blastocoel) | scRNAseq_E5_2 |
| 2501  Large White | 27/01/2021 | SAMEA112465592 | 2000966  crossed | normal breeding procedure | No | E5 | 9 | embryo_D5_5  embryo_single_cell_D5_3 | SAMEA112465624  SAMEA112465659 | early blastocyst (compact) | scRNAseq_E5_1 |
| 2403  Large White | 27/01/2021 | SAMEA112465593 | 2000966  crossed | normal breeding procedure | No | E5 | 10 | embryo_D5_6  embryo_single_cell_D5_7 | SAMEA112465625  SAMEA112465663 | early blastocyst (with blastocoel) | scRNAseq_E5_2 |
| 2330  Large White | 27/01/2021 | SAMEA112465594 | 2000966  crossed | normal breeding procedure | No | E5 | 6 | embryo_D5_7  embryo_single_cell_D5_4 | SAMEA112465626  SAMEA112465660 | early blastocyst (compact) | scRNAseq_E5_1 |
| 4290  Large White | 19/05/2021 | SAMEA112465595 | 1245  Large White | normal breeding procedure | Yes  4290_E7-4 | E7 | 6 | embryo_D7_7  embryo_single_cell_D7_4 | SAMEA112465629  SAMEA112465664 | spheroid blastocyst | scRNAseq_E7_3 |
| 4239  Large White | 19/05/2021 | SAMEA112465596 | 1245  Large White | normal breeding procedure | Yes  4239_E7-3 | E7 | 5 | embryo_D7_9  embryo_single_cell_D7_5 | SAMEA112465631  SAMEA112465665 | hatched blastocyst | scRNAseq_E7_4 |
| 4220  Large White | 19/05/2021 | SAMEA112465597 | 1245  Large White | normal breeding procedure | No | E7 | 7 | embryo_D7_11  embryo_single_cell_D7_6 | SAMEA112465633  SAMEA112465666 | hatched blastocyst | scRNAseq_E7_4 |
| 4182  Large White | 19/05/2021 | SAMEA112465599 | 1245  Large White | normal breeding procedure | Yes  4182_E7-2 | E7 | 5 | embryo_D7_13  embryo_single_cell_D7_7 | SAMEA112465635  SAMEA112465667 | hatched blastocyst | scRNAseq_E7_4 |
| 4289  Large White | 19/05/2021 | SAMEA112465600 | 1245  Large White | normal breeding procedure | No | E7 | 4 | embryo_D7_15  embryo_single_cell_D7_8 | SAMEA112465637  SAMEA112465668 | hatched blastocyst | scRNAseq_E7_4 |
| 4246  Large White | 19/05/2021 | SAMEA112465601 | 1245  Large White | normal breeding procedure | No | E7 | 5 | embryo_D7_17  embryo_single_cell_D7_9 | SAMEA112465639  SAMEA112465669 | hatched blastocyst | scRNAseq_E7_4 |
| 4181  Large White | 19/05/2021 | SAMEA112465602 | 1245  Large White | normal breeding procedure | Yes  4181_E7-1 | E7 | 4 | embryo_D7_19  embryo_single_cell_D7_10 | SAMEA112465641  SAMEA112465670 | hatched blastocyst | scRNAseq_E7_4 |
| 4088  Large White | 03/06/2021 | SAMEA112465603 | 1000  Large White | normal breeding procedure | Yes  4088_E9-1 | E9 | 7 | embryo_D9_6  embryo_single_cell_D9_3 | SAMEA112465643  SAMEA112465671 | small ovoid blastocyst | scRNAseq_E9_3 |
| 4278  Large White | 03/06/2021 | SAMEA112465604 | 1000  Large White | normal breeding procedure | Yes  4278_E9-2 | E9 | 3 | embryo_D9_7  embryo_single_cell_D9_4 | SAMEA112465644  SAMEA112465672 | ovoid blastocyst 5mm | scRNAseq_E9_4 |
| 4300  Large White | 03/06/2021 | SAMEA112465605 | 1000  Large White | normal breeding procedure | Yes  4300_E9-3 | E9 | 3 | embryo_D9_9  embryo_single_cell_D9_5 | SAMEA112465646  SAMEA112465673 | ovoid blastocyst 5mm | scRNAseq_E9_4 |
| 4221  Large White | 03/06/2021 | SAMEA112465606 | 1000  Large White | normal breeding procedure | Yes  4221_E11-3 | E11 | 2 | embryo_D11_4  embryo_single_cell_D11_1 | SAMEA112465648  SAMEA112465674 | ovoid blastocyst 10-15mm | scRNAseq_E11_1 |
| 4086  Large White | 03/06/2021 | SAMEA112465607 | 1000  Large White | normal breeding procedure | Yes  4086_E11-2 | E11 | 2 | embryo_D11_6 embryo_single_cell_D11_2 | SAMEA112465650  SAMEA112465675 | ovoid blastocyst 10-15mm | scRNAseq_E11_2 |
| 2548  Large White | 02/02/2021 | SAMEA112465595 | 2000966  crossed | Normal breeding procedure | Yes  2548_E11-1 | E11 | 10 |  |  |  |  |
| 1384  Large White | 02/02/2021 |  | 1111  crossed | Normal breeding procedure | Yes  1384_E5-1 | E5 | 10 |  |  |  |  |
| 1982  Large White | 03/02/2021 |  | 610  crossed | Normal breeding procedure | Yes  1982_E5-2 | E5 | 12 |  |  |  |  |
| 2044  Large White | 03/02/2021 |  | 610  crossed | Normal breeding procedure | Yes  2044_E5-3 | E5 | 14 |  |  |  |  |
| 1844  Large White | 02/02/2021 |  | 1111  crossed | Normal breeding procedure | Yes  1844_E5-5 | E5 | 13 |  |  |  |  |
| 2048  Large White | 03/02/2021 |  | 610  crossed | Normal breeding procedure | Yes  2048_E5-6 | E5 | 10 |  |  |  |  |
