## Supplementary Table 2 for "Cell specification and functional interactions in the pig blastocyst inferred from single cell transcriptomics"

| **Samples** | **Number of Reads** | **Reads Mapped Confidently to Exonic Regions** | **Median UMIs Counts per Cell** | **Number of genes** | **Number of cells before quality controls** | **Minimal number of UMIs per cells** | **Maximal number of features per cells** | **Maximal % of mitochondrial RNA per cells** | **Number of cells after quality controls** | **Number of principal composant used** | **Clustering resolution** |
| --- | --- | --- | --- | --- | --- | --- | --- | --- | --- | --- | --- |
| E5-1 | 303246545 | 75.1 | 762 | 18627 | 1831 | 500 | 10000 | 25 | 919 | 15 | 0.1 |
| E5-2 | 328087638 | 72.2 | 1502 | 21603 | 980 |  |  |  | 307 |  |  |
| E7-1 | 216372505 | 80.2 | 1358 | 12903 | 1249 | 300 |  | 10 | 1171 | 20 | 0.05 |
| E7-2 | 152525069 | 73.5 | 1232 | 13529 | 1212 |  |  |  | 962 |  |  |
| E7-3 | 266879834 | 59.9 | 2.937 | 23900 | 1492 |  |  |  | 767 |  |  |
| E7-4 | 220003973 | 55.8 | 1626 | 24297 | 2124 |  |  |  | 1328 |  |  |
| E9-1 | 157253423 | 77.8 | 984 | 15972 | 1286 | 500 |  |  | 581 | 20 | 0.1 |
| E9-2 | 144826623 | 79.0 | 1063 | 17801 | 4798 |  |  |  | 1969 |  |  |
| E9-3 | 245106853 | 66.7 | 1613 | 23553 | 4494 |  |  |  | 2827 |  |  |
| E9-4 | 211348253 | 67.3 | 1243 | 22941 | 9940 |  |  |  | 7350 |  |  |
| E11-1 | 225484028 | 67.6 | 1085 | 21399 | 9889 | 300 |  | 20 | 9474 | 20 | 0.1 |
| E11-2 | 221048966 | 67.7 | 1207 | 20675 | 7650 |  |  |  | 7233 |  |  |
| Tissue |  | | | | | | | | | | |
| TE |  | | | | | | | | | 25 | 0.2 |
| HYPO |  |  |  |  |  |  |  |  |  | 20 | 0.1 |
